# TFEB degradation is regulated by an IKK/β-TrCP2 phosphorylation-ubiquitination cascade

**DOI:** 10.1101/2023.10.16.562572

**Authors:** Yan Xiong, Jaiprakash Sharma, Meggie N. Young, Wen Xiong, Ali Jazayeri, Karl F. Poncha, Ma. Xenia G. Ilagan, Qing Wang, Hui Zheng, Nicolas L. Young, Marco Sardiello

## Abstract

Transcription factor EB (TFEB) is a master regulator of lysosomal biogenesis and autophagy that plays a key role in the regulation of cellular clearance pathways. TFEB is regulated via a complex array of post-translational modifications (PTMs), but the exact molecular mechanism that regulates TFEB stability has remained elusive. Here, we show that TFEB levels are critically regulated by a defined phosphorylation-ubiquitination cascade. A human kinome screen identifies IKK (inhibitor of κB kinase) as a TFEB modifier, and a combination of phosphorylation assays, mass spectrometry analyses, and site-specific mutagenesis unveils a previously unrecognized TFEB phospho-degron (^423^SPFPSLS^429^) as the target of IKK. We show that IKK-mediated phosphorylation of TFEB triggers ubiquitination of adjacent lysine residues (K430 and K431) by the E3 ligase β-TrCP2 (β-Transducin repeat-containing protein 2), thereby tagging TFEB for degradation. Modified TFEB constructs that abolish these PTMs show much increased stability and expression levels but remain equally sensitive to autophagy- or stress-related stimuli while maintaining the capability to promote the expression of TFEB target genes and the clearance of Alzheimer’s associated tau in a cellular model of disease. Our results therefore uncover an IKK/β-TrCP2 phosphorylation-ubiquitination cascade as a major mechanism that governs TFEB stability independently of other TFEB regulators.

## INTRODUCTION

Transcription factor EB (TFEB) is a master regulator of the autophagy-lysosomal pathway (ALP)^1^. Active TFEB promotes the transcription of lysosomal and autophagy genes^1–3^ and participates in the induction of ALP-based pathways such as lysosome-mediated clearance^1^, lysosomal proteostasis^4^, and lysosomal exocytosis^5^. TFEB also participates in additional cellular pathways that include the ER stress response^4, 6^ and mitochondrial biogenesis^7, 8^. TFEB plays a pivotal role in the regulation of lysosome-to-nucleus communication by translating inputs from mechanistic target of rapamycin complex 1 (mTORC1)^9–13^ and other key metabolic regulators including protein kinase B (PKB/AKT)^14, 15^ and adenosine monophosphate-activated protein kinase (AMPK)^16^, all of which have specific roles at the lysosome^17–19^. Importantly, these and other kinases modify TFEB on distinct subsets of amino acids^9–16, 20–23^, indicating that TFEB has several independent layers of regulation that presumably execute different branches of the cellular adaptive response or serve additional functions. Studies with cultured cells have shown that, in normal cell growth conditions, TFEB protein is predominantly found in the cytosol in an inactive state. TFEB cytosolic localization is facilitated by the binding of YWHA/14-3-3 proteins to TFEB phospho-Ser211, which sequesters TFEB within the cytosolic compartment^12, 13^. Ser211 is phosphorylated by active mTORC1^12, 13^, which therefore limits the ALP-enhancing functions of TFEB. Conversely, lysosomal storage, cell starvation, and other stressors promote TFEB nuclear translocation^1, 2, 6, 11–13, 24^. In the nucleus, TFEB binds to the “coordinated lysosomal expression and regulation” (CLEAR) motif located within the promoter regions of genes participating in lysosomal pathways and promotes their transcription^1^. Therefore, TFEB nuclear translocation is regarded as an indicator of TFEB activation.

Active TFEB and ALP enhancement promote cellular clearance, aiding in the elimination of protein aggregates and lysosomal storage material. By leveraging this principle, we and others have shown that TFEB is both part of pathogenic cascades and a candidate therapeutic target in neurodegenerative storage disorders and beyond^1, 4, 5, 14, 25–30^. Multiple studies have demonstrated that exogenous TFEB expression reduces aggregated tau and β-amyloid in cell lines as well as in brain tissues of mouse models of Alzheimer’s disease^26, 28, 31^, highlighting the significance of TFEB as a potential target for therapeutic intervention.

While best known for their roles in Nuclear Factor kappa B (NF-κB) signaling, the components of the IκB kinase (IKK) complex have more recently emerged as NF-κB-independent regulators of processes as diverse as mTORC1 signaling^32–34^, apoptosis^35, 36^, cell proliferation and differentiation^37, 38^, and epigenetic control of gene transcription^39–41^ through direct phosphorylation of key players of these processes. The IKK complex is composed of three components: the partially redundant catalytic subunits IKKα (encoded by the *CHUK* gene) and IKKβ (*IKBKB*), and a scaffold/regulatory subunit, IKKγ (*NEMO*)^42–44^. IKK has been connected with the mTOR pathway through two separate but converging mechanisms. IKK was first shown to be an inhibitory modifier of Tuberous Sclerosis Complex 1 (TSC1)^32^, a key negative regulator of mTORC1 signaling^45^. Follow-up work has shown that IKK subunits also associate with and phosphorylate mTOR, an event that promotes mTORC1 kinase activity^33, 34^. Thus, active IKK functions as a positive regulator of mTORC1 according to at least two different mechanisms. Activation of IKK can occur through various stimuli, including TNFα signaling^43^. For some of its protein targets, phosphorylation by IKK generates a phosphodegron, a specific signal that is recognized by the beta-transducin repeats-containing proteins (β-TrCP)—F-box proteins that serve as the substrate recognition subunits for the SCF^β-TrCP^ E3 ubiquitin ligases^46–49^. The mammalian members of the β-TrCP family, the β-TrCP1 and β-TrCP2 paralogs, share sequence similarity and biochemical properties but may have distinct substrates^46^. For example, β-TrCP2 has been characterized as the dominant paralog in the regulation of autophagy and cell growth via starvation-induced recognition and ubiquitination of β-TrCP1 and the mTORC1 inhibitors, DEPTOR and REDD1^50^.

In this study, we show that TFEB protein levels in normal cell growth conditions are markedly regulated by a phosphorylation-ubiquitination cascade executed by IKK and β-TrCP2. We find that IKK phosphorylates TFEB at Ser427, a previously unrecognized modification site of TFEB, and other surrounding serine residues. IKK phosphorylation of TFEB triggers the generation of a phosphodegron that is recognized by β-TrCP2, which then promotes degradative ubiquitination of TFEB at lysine sites adjacent to the phosphodegron. Selective genetic disruption of this phosphorylation-ubiquitination cascade increases TFEB protein levels by several fold but does not intersect with the signaling controlling TFEB nuclear translocation or TFEB’s capability to enhance transcription of its target genes and cellular clearance. This previously unrecognized mechanism clarifies mechanistically how TFEB is targeted for proteasomal degradation and introduces a novel and powerful layer of regulation of TFEB that warrants future exploration as a candidate therapeutic target in diseases with impaired cellular clearance.

## RESULTS

### IKK complex regulates TFEB stability

To efficiently screen for kinases that modify TFEB levels post-translationally, we generated a TFEB-GFP fusion construct as a reporter readout that does not depend on transcriptional regulation of gene levels and does not require incubation with antibodies. We first tested this construct by transient transfection in HeLa cells by using fluorescence microscopy to monitor TFEB protein levels and subcellular localization. Calculation of a TFEB nuclear translocation index (TFEBNTI, see Methods for details) for hundreds of cells showed that TFEB nuclear localization levels are positively associated with TFEB protein levels (Supplementary Figure 1a, b), presumably because high levels of TFEB protein saturate the systems that promote TFEB cytosolic retention. Thus, we generated cells stably expressing TFEB-GFP (henceforth referred to as HeLa/TFEB-GFP) by selecting a non-clonal pool of cells infected with a TFEB-GFP lentivirus. Fluorescence microscopy analysis showed that TFEB was generally expressed at lower levels compared to transiently transfected cells, and that the levels of TFEB nuclear localization were independent of TFEB expression levels in HeLa/TFEB-GFP cells (Supplementary Figure 1c, d), indicating the suitability of these cells for screening procedures.

To identify candidate kinases that regulate TFEB stability, we carried out a human kinome screen using a library containing 436 established kinase inhibitors. For each inhibitor, we applied five different combinations of incubation time and drug concentration, with cells incubated with either 1 or 3 µM kinase inhibitors for 1 hour, or cells incubated with 1, 3, or 10 µM kinase inhibitors for 6 hours. We used automated fluorescence microscopy to acquire images of hundreds to thousands of cells per condition and assess TFEB-GFP signal levels in whole cell as well as in the cytosol and nuclei separately. In each plate, Torin1 (a strong catalytic inhibitor of mTORC1) was used as a positive control for its ability to modify cytosolic and nuclear TFEB levels, while DMSO was used as a vehicle control. To assess the robustness of the pipeline, we integrated the microscopy data from the five conditions to perform unsupervised hierarchical clustering of the drugs based on their effect on TFEB nuclear translocation (Table S1). The analysis showed that the rapalogs (allosteric mTORC1 inhibitors) present in the library clustered with Torin1 as expected (Fig. 1a, red arrow), thereby validating the pipeline. The vast majority of the kinase inhibitors screened clustered with DMSO, indicating little to no effects on TFEB subcellular distribution. Interestingly, a small cluster of kinase inhibitors that increased TFEB nuclear signal was distinct from the cluster of mTORC1 inhibitors (indicated by the green arrow in Fig. 1a), suggesting the occurrence of additional mechanisms contributing to the change in cytosolic and/or nuclear TFEB levels. A scatterplot of total TFEB-GFP intensity versus the product of the distances from Torin1 and vehicle in the hierarchical cluster highlighted a small group of kinases whose inhibition simultaneously increased TFEB nuclear intensity and total TFEB intensity—IKK, Axl, JAK2, and CDK4/6 (Fig. 1b). CDK4/6 is a known kinase modifier of TFEB^23^, and both Axl and JAK2 are general receptor tyrosine kinases that modulate the PI3K/Akt/mTOR pathway among many others^51, 52^. IKK has multiple strong connections with mTORC1 signaling^32–34^ but has never been studied in the context of TFEB biology. Thus, we selected to focus on IKK for functional studies.

**Fig. 1.**
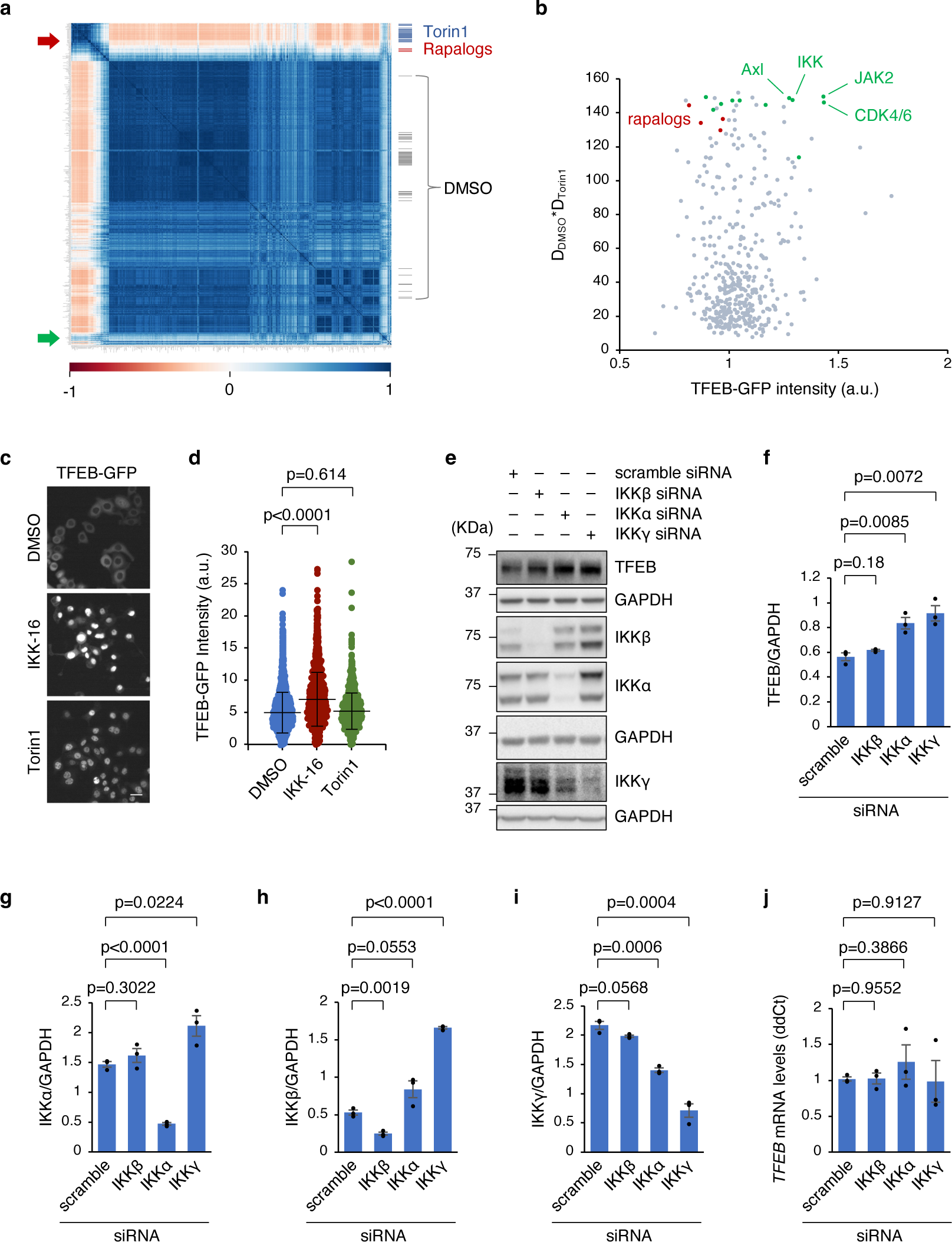
A human kinome screen identifies IKK as a candidate TFEB modifier. **a** Hierarchical clustering of all tested drugs based on TFEBNTI distribution curves. The red arrow indicates a cluster of drugs targeting mTORC1, while the green arrow indicates a cluster of drugs affecting TFEB subcellular distribution, but distinct from the mTORC1 cluster. Vehicle controls (DMSO) are indicated in the clustering. **b** Dot plot displaying TFEB-GFP intensity levels and the product of the distances from DMSO and Torin1 in the hierarchical clustering. TFEB-GFP intensity levels were measured after 6-hour incubation with a drug concentration of 10 µM. **c** Representative images of the kinome screen showing HeLa/TFEB-GFP treated with DMSO, IKK-16, or Torin1. Scale bar: 30 μm. **d** Quantification of mean TFEB intensity of cells from the kinome screen under the above-mentioned conditions. Results are presented as mean ± SD. n > 900 cells per condition. **e** Immunoblot analysis of HEK-293T cells transiently transfected with scramble, *IKKα*, *IKKβ*, or *IKKγ* siRNA (n = 3 independent experiments) using the indicated antibodies. **f-i** Quantification of the immunoblots in (**e**), normalized to corresponding GAPDH levels. **j** Relative *TFEB* mRNA levels determined by RT-qPCR in samples shown in panel (**e**). The expression levels were normalized to the housekeeping gene, *GAPDH*. Results in (**f-j**) are presented as mean ± S.E.M. *p* values are calculated based on the Student’s *t*-test.

The observed changes in TFEB subcellular localization that follows IKK inhibition are in line with the notion that active IKK promotes the activity of mTORC1^32–34^, the best characterized inhibitor of TFEB nuclear translocation^9–13^. Indeed, we confirmed that IKK inhibition in the conditions used in the screen resulted in mTORC1 inhibition (Supplementary Figure 2a, b) and TFEB nuclear translocation (Fig. 1c). However, IKK inhibition also resulted in a strong increase in TFEB signal intensity, whereas direct catalytic inhibition of mTORC1 by Torin1 did not (Fig. 1c, d), indicating that the two processes are uncoupled.

We next tested whether endogenous TFEB levels are regulated by IKK by monitoring TFEB protein levels in HEK293T cells upon siRNA-mediated silencing of either IKK component. Immunoblot analyses showed that silencing either *CHUK*/IKKα or *NEMO*/IKKγ increased TFEB protein levels (Fig. 1e, f), indicating that IKK does play a role in TFEB regulation in normal cell growth conditions. IKKγ silencing also increased the levels of IKKα and IKKβ (Fig. 1g, h), whereas IKKα silencing decreased IKKγ levels (Fig. 1i), suggesting the presence of feedback or regulatory loops that mutually influence protein stability amongst the IKK components. An RT-qPCR analysis showed that *TFEB* mRNA levels were not affected by silencing of either IKK component (Fig. 1j). Taken together, these data indicate that TFEB levels may be dependent on the overall activity of the IKK complex, and that IKK regulation of TFEB is post-transcriptional.

### IKK phosphorylates TFEB at multiple serine residues

IKKα and IKKβ are structurally similar but only partially redundant in function, as shown by their selectivity for their phosphorylation substrates^44, 53, 54^. We tested whether IKKα and IKKβ phosphorylate TFEB by performing phos-tag analyses. In phos-tag analyses, the rate of protein migration depends on the number of phosphorylated residues, and multiple bands indicate the presence of multiple phosphorylated sites on the same protein^55^. 3xFlag-tagged TFEB was transiently expressed in HeLa cells, immunoprecipitated, and dephosphorylated *in vitro* by *λ* phosphatase before being mixed with commercially available IKKα or IKKβ for phos-tag analysis. The results showed that both IKKα and IKKβ phosphorylated TFEB, with IKKβ showing higher efficacy and producing multiple phosphorylated bands (Fig. 2a). In a control reaction, *in vitro* phosphorylation of TFEB by Akt using the same procedure resulted in a single phosphorylated TFEB band which was completely abolished by alanine substitution of serine 467, as expected^14^ (Supplementary Fig. 3a). Thus, the presence of multiple phosphorylated bands after incubation with IKKβ indicates the presence of multiple phosphorylation events mediated by IKKβ.

**Fig. 2.**
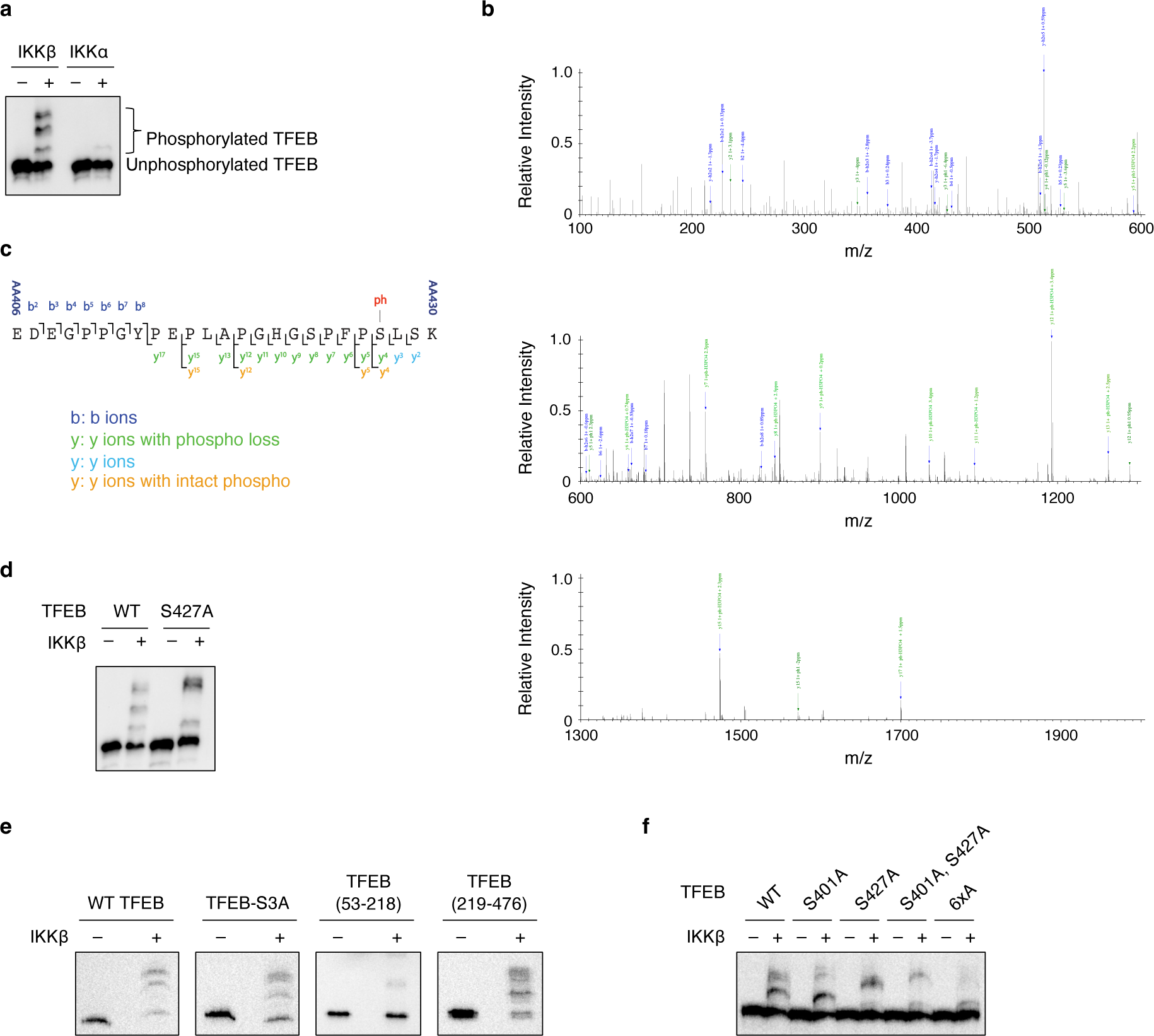
IKK phosphorylates TFEB. **a** TFEB-3xFlag was immunoprecipitated from HeLa cell lysate, dephosphorylated by λ phosphatase, and subjected to a kinase assay with recombinant IKKβ or IKKα. The reaction was stopped by 2x Laemmli buffer and analyzed using phos-tag analysis with a TFEB antibody. **b** Higher energy collisional dissociation (HCD) MS2 mass spectrum of TFEB’s tryptic peptide from precursor *m/z* 881.73 corresponding to the amino acid sequence 406AA-430AA (EDEGPPGYPEPLAPGHGSPFPSLSK) generated by the Fusion Lumos Orbitrap Mass Spectrometer at mass range from 100 to 600 *m/z* (*upper panel*), mass range from 600 to 1300 *m/z* (*middle panel*), and mass range from 1300 to 2000 *m/z* (*lower panel*). **c** Summary map of all MS2 fragment ions that identify the TFEB sequence 406AA- 430AA and localize the phosphorylation to S427. **d** *In vitro* kinase assays. Immunoprecipitated TFEB-3xFlag or TFEB S427A-3xFlag were incubated with recombinant IKKβ and analyzed as described in (**a**). **e** *In vitro* kinase assays. Immunoprecipitated TFEB-3xFlag or TFEB S3A- 3xFlag or 3xFlag-tagged TFEB fragments were incubated with recombinant IKKβ and analyzed as described in (**a**). **f** *In vitro* kinase assays. Immunoprecipitated TFEB-3xFlag or 3xFlag-tagged TFEB mutants with specific amino acid substitutions were incubated with recombinant IKKβ and analyzed as described in (**a**).

*In vitro* peptide screens to define IKKα/β’s substate preferences, along with the identification of several bona fide substrates, have shown that IKKα and IKKβ do not have strict requirements for the amino acids surrounding their target serine, which makes predicting their candidate substrate sites difficult^54, 56, 57^. To identify the TFEB residues phosphorylated by IKK, we used mass spectrometry to analyze TFEB upon *in vitro* phosphorylation by IKKβ. The results showed abundant phosphorylation localized to serine 427 by multiple fragment ions, including fragment ions with intact phosphoserine (Fig. 2b, c). Specifically, the data showed the presence of an intact phosphorylated peptide at 406AA-430AA with *m/z* 881.7311 (-1.99 ppm error, expected mass of 881.732851 *m/z*) (Supplementary Fig. 3b). MS2 analysis (in which precursor peptide ions are fragmented and analyzed to derive sequence information, including PTM localization) resulted in multiple ion types supporting phosphorylation localization at S427. Similar MS2 results were observed for the phosphorylated 406AA-431AA peptide that arises from incomplete digestion due to the ‘KK’ motif at 430-431AA (not shown). In phos-tag analyses, alanine substitution of S427 changed the pattern of phospho-TFEB bands and reduced phosphorylation efficiency of TFEB by IKKβ, resulting in higher levels of unphosphorylated TFEB-S427A compared to wild-type (WT) TFEB (Fig. 2d). To narrow down the additional sites targeted by IKKβ, we first tested S3, the only serine present within the N-terminal domain of TFEB, and found that alanine substitution did not change the TFEB phosphorylation pattern (Fig. 2e). Therefore, we excluded S3 as a candidate site. We then separately analyzed the phosphorylation of TFEB’s 53-218 and 219-476 fragments (Supplementary Fig. 3c). The results showed that TFEB (53-218) was not phosphorylated by IKKβ, while TFEB (219-476) presented multiple phosphorylation bands (Fig. 2e). These results indicate that the additional target sites of IKK are also located within the 219-476 region. To identify these sites, we generated a library of TFEB mutants with single alanine substitutions for each of the 24 serine residues present within the 219-476 region and evaluated the phosphorylation status of each mutant by *in vitro* kinase assay with IKKβ. The results showed that alanine substitution of S401 and S429 also affected the phosphorylation pattern of TFEB (Supplementary Fig. 3d). The S401A/S427A double mutant showed a significant reduction in phosphorylation efficiency compared to WT TFEB, although a small fraction of phosphorylated protein was still observed (Fig. 2f). Alanine substitution of four additional adjacent serine residues (S397, S399, S423, and S429) along with S401 and S427 (henceforth referred to as TFEB-6xA) thoroughly abolished TFEB phosphorylation by IKKβ (Fig. 2f). Interestingly, these six sites have the appearance of two somewhat similar modules each the approximate size of a typical kinase interaction surface, S397/399/401 (FSHSLSF) and S423/427/429 (SPFPSLS). We have shown that S427 is the predominant site of IKKβ-mediated phosphorylation for SPFPSLS. Similarly, our results suggest that S401 is likely the predominant site of phosphorylation for FSHSLSF. Mutations of residues in such proximity can act indirectly; however, residual phosphorylation indeed appears to occur at neighboring serines at least in these mutants with unknown effect. Thus, we next used TFEB-6xA to completely abolish all IKKβ activity on TFEB to isolate this novel pathway for functional analysis.

### IKK phosphorylation-resistant TFEB displays increased expression levels and stability

To investigate the functional consequences of abolishing IKK-mediated TFEB phosphorylation on TFEB stability, we generated non-clonal pools of HeLa cells stably expressing 3xFlag-tagged WT TFEB or TFEB-6xA (henceforth referred to as HeLa/TFEB and HeLa/TFEB-6xA). Immunoblot analyses showed that, in normal cell growth condition, TFEB-6xA protein had much higher levels than WT TFEB (Fig. 3a, b), despite no significant differences between the RNA expression levels of the two constructs (Fig. 3c). To test whether this effect is caused by differences in protein stability, we used cycloheximide to block protein synthesis and measured degradation rates for either TFEB construct. The results showed that the levels of WT TFEB rapidly decreased upon blockage of protein synthesis, whereas the levels of TFEB-6xA remained stable within the monitored time frame (Fig. 3d). These data indicate that abolishing IKK phosphorylation sites increases the stability of TFEB. We next tested the stability of WT and 6xA TFEB under TNFα-induced activation of IKK. The results showed reduction of WT TFEB levels by 30 min of TNFα treatment, whereas the levels of TFEB-6xA remained unchanged (Fig. 3e). Previous work has shown that TFEB levels are reduced by treatment with lipopolysaccharide (LPS), a stress inducer that causes the activation of inflammatory responses in various cell types^21, 58^. Immunoblot analyses showed that LPS treatment indeed reduced the levels of WT TFEB in a dose-dependent manner; however, it had a lesser impact on TFEB-6xA levels (Fig. 3f). In summary, our findings demonstrate that the abolishment of IKK target sites increases TFEB protein expression levels and stability, indicating that IKK phosphorylation of TFEB promotes TFEB degradation.

**Fig. 3.**
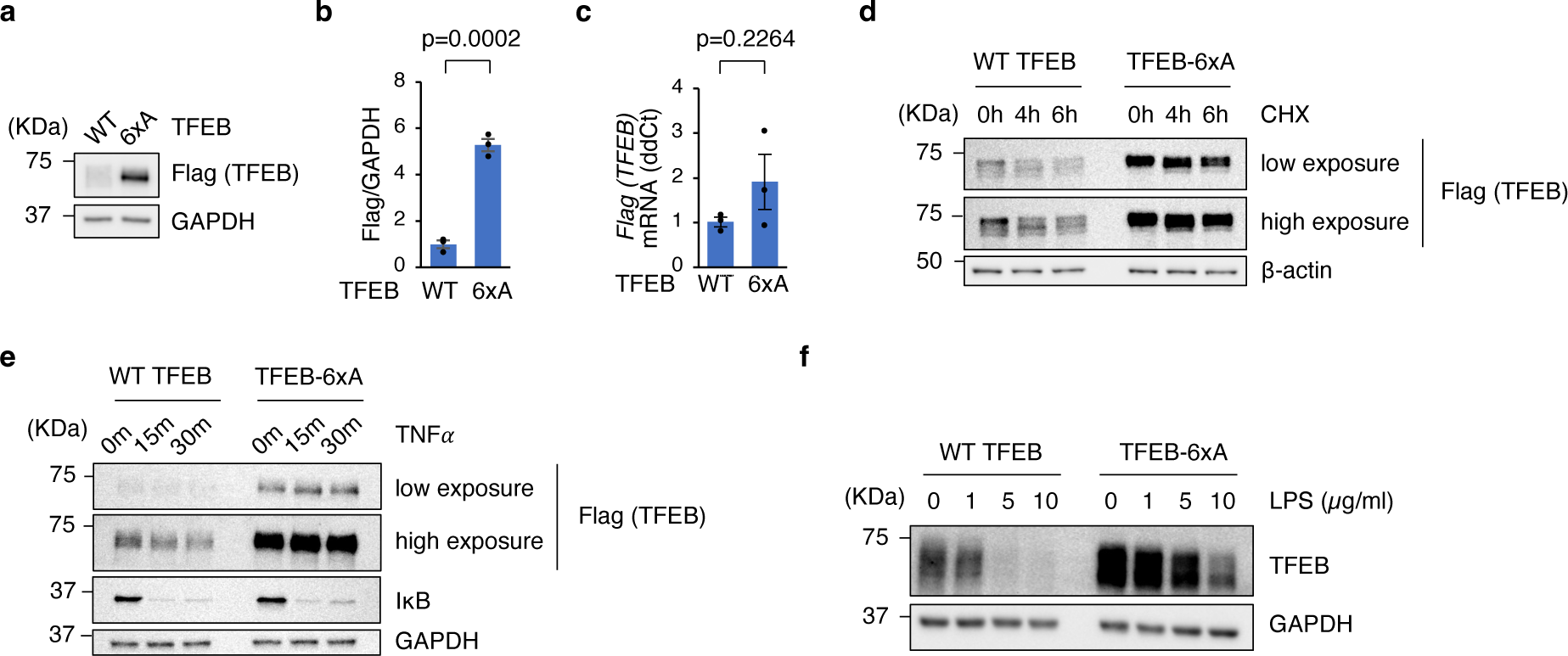
Abolishment of IKK phosphorylation sites stabilizes TFEB. **a** Immunoblot analysis of HeLa/TFEB-3xFlag and HeLa/TFEB-6xA-3xFlag using the indicated antibodies (n = 3 independent replicates for each group). **b** Quantification of the immunoblots in (**a**), normalized to corresponding GAPDH levels. **c** Relative *TFEB-3xFlag* mRNA levels determined by RT- qPCR in samples shown in panel (**a**). The expression levels were normalized to the housekeeping gene, *GAPDH*. Results in (**b**) and (**c**) are presented as mean ± S.E.M. *p* values are calculated based on the student’s *t*-test. **d** HeLa/TFEB-3xFlag and HeLa/TFEB-6xA-3xFlag were treated with 10 μg/ml cycloheximide (CHX), and the cells were harvested at indicated time points. Immunoblot analyses were performed using the indicated antibodies. **e** HeLa/TFEB-3xFlag and HeLa/TFEB-6xA-3xFlag were treated with 20 ng/ml TNFα, and the cells were harvested at indicated time points. Immunoblot analyses were performed using the indicated antibodies. **f** HeLa/TFEB-3xFlag and HeLa/TFEB-6xA-3xFlag were treated with LPS at the specified concentrations for 48 h, and the cells were harvested for immunoblot analyses using the indicated antibodies.

### β-TrCP2 mediates TFEB ubiquitination and stability downstream of IKK phosphorylation

IKK-mediated phosphorylation primes certain protein substrates for degradative ubiquitination by generating a phosphodegron that is recognized by the E3 ubiquitin ligases β-TrCP1/2^47–49^. To investigate whether β-TrCPs mediate TFEB ubiquitination, we assessed TFEB ubiquitination levels upon exogenous expression of β-TrCP1, β-TrCP2, or the E3 ubiquitin ligases STUB1 (STIP1 homology and U-Box containing protein 1) and DCAF7 (DDB1 and CUL4 associated factor 7), which had previously been associated with TFEB degradation^59, 60^. Coexpression with HA-tagged ubiquitin and TFEB-3xFlag followed by TFEB immunoprecipitation and HA immunoblotting showed that, in normal cell growth conditions, β-TrCP2—but not β-TrCP1— promoted TFEB ubiquitination (Fig. 4a). Next, we silenced *β-TrCP2* to test whether β-TrCP2 mediates TFEB ubiquitination. Blockage of proteasomal degradation by low concentrations of carfilzomib (CFZ) resulted in the accumulation of ubiquitinated TFEB, which was completely prevented by *β-TrCP2* silencing (Fig. 4b). These data show that β-TrCP2 functions as a ‘default’ TFEB ubiquitin ligase and indicate that TFEB tagging for proteasomal degradation is a continuously occurring process in normal cell growth conditions. Due to the unavailability of suitable antibodies against endogenous β-TrCP2, the efficiency of β-TrCP2 silencing was evaluated by monitoring the levels of endogenous *β-TrCP2* mRNA (Supplementary Fig. 4a) and the protein levels of a HA-β-TrCP2 construct transiently transfected into HEK293T cells (Supplementary Fig. 4b). Next, we tested whether IKK-mediated phosphorylation of TFEB is necessary for TFEB ubiquitination by β-TrCP2. To this aim, we performed sequential transfections in HEK293T cells by first introducing either WT TFEB or TFEB-6xA constructs along with HA-tagged ubiquitin, and subsequently transfecting the cells with siRNAs against β- TrCP2 or scrambled siRNAs. Immunoprecipitation of TFEB followed by HA immunoblotting showed that IKK-resistant TFEB-6xA exhibits low levels of ubiquitination compared to WT TFEB (Fig. 4c). *β-TrCP2* silencing abated the ubiquitination levels of WT TFEB as expected but did not modify those of TFEB-6xA (Fig. 4c and Supplementary Fig. 4c). Remarkably, scramble siRNA-treated TFEB-6xA displayed ubiquitination levels similar to WT TFEB upon *β-TrCP2* silencing, indicating that elimination of IKK’s target sites from TFEB mimics a lack of engagement of β-TrCP2.

**Fig. 4.**
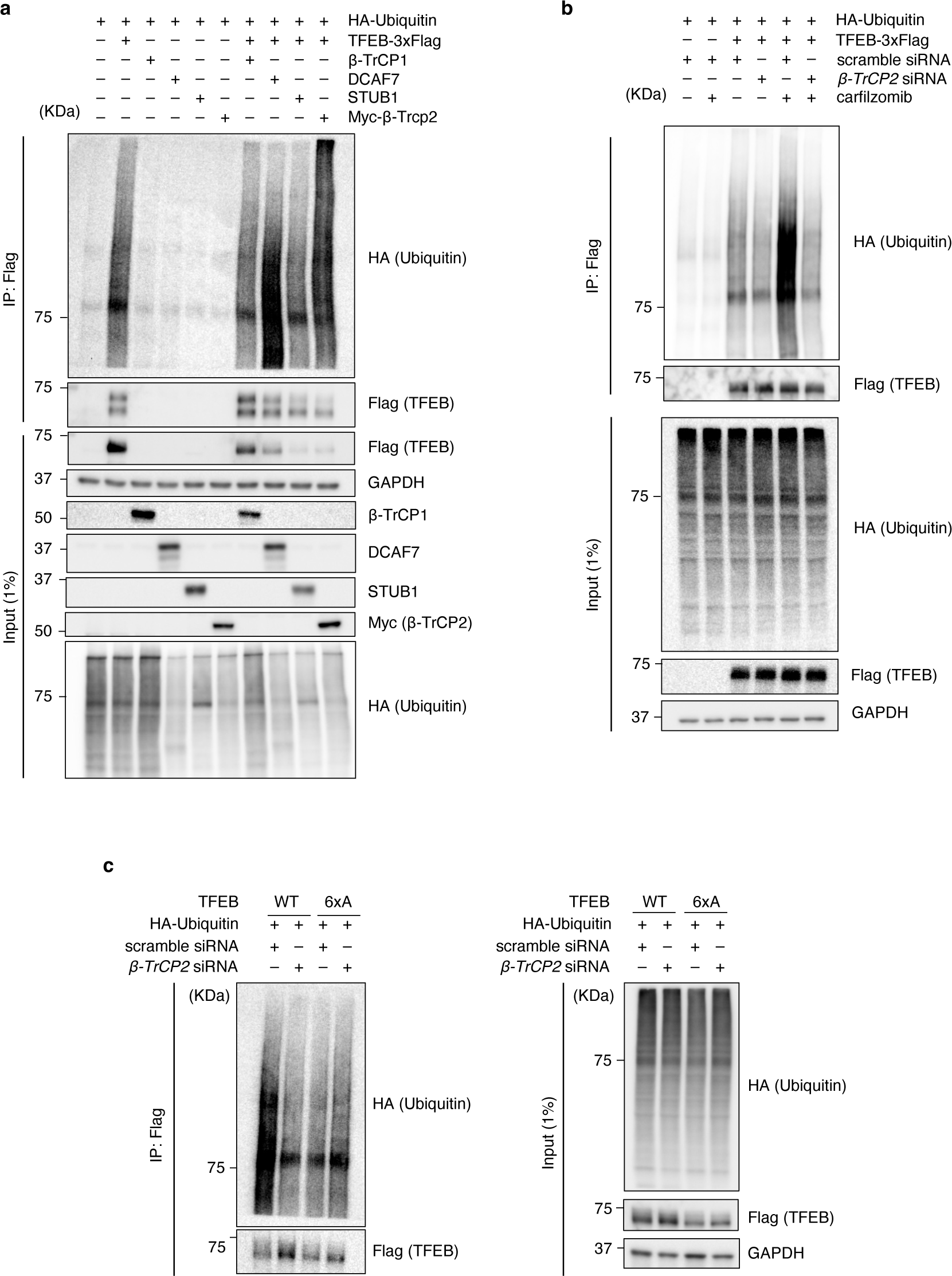
β-TrCP2 mediates TFEB ubiquitination downstream of IKK phosphorylation. **A** HEK-293T cells were cotransfected with TFEB-3xFlag, HA- ubiquitin, and one of the indicated E3 ligases (β-Trcp1/2, STUB1 or DCAF7). 24 h after transfection, TFEB proteins were immunoprecipitated and the ubiquitination levels were checked using the HA antibody. **b** HEK- 293T cells were cotransfected with HA-ubiquitin and TFEB-3xFlag. 24 h after transfection, the cells were subjected to a second round of transfection with scramble or *β-Trcp2* siRNA. After an additional 28 hrs, the cells were incubated with 100 nM CFZ for 20 h before harvest. TFEB proteins were then immunoprecipitated, and ubiquitination levels were assessed using an HA antibody. **c** HEK-293T cells were cotransfected with HA-ubiquitin and TFEB-3xFlag or TFEB- 6xA-3xFlag. 24 h after transfection, the cells were subjected to a second round of transfection with scramble or *β-Trcp2* siRNA. After an additional 48 hrs, TFEB proteins (WT and 6xA) were immunoprecipitated, and ubiquitination levels were assessed using an HA antibody.

*β-TrCP2* silencing in HEK293T cells resulted in a significant increase of endogenous TFEB levels (Fig. 5a-c), confirming that β-TrCP2 participates in TFEB turnover in normal cell growth conditions. In contrast to TFEB protein levels, the mRNA levels of TFEB remained unchanged upon β-TrCP2 silencing (Fig. 5d), confirming that β-TrCP2 affects TFEB protein levels post-transcriptionally. In addition, we investigated mechanistically whether β-TrCP2-mediated modulation of TFEB levels requires the IKK-targeted phosphodegron by testing the effects of *β- TrCP2* silencing on the IKK-resistant TFEB-6xA construct. To enable a comparison with WT TFEB, we silenced β-TrCP2 in both HeLa/TFEB and HeLa/TFEB-6xA. The results showed that *β-TrCP2* silencing almost doubled the levels of WT TFEB in normal cell growth conditions, while it had a minimal effect on the levels of TFEB-6xA (Fig. 5e-g). Collectively, these findings demonstrate that β-TrCP2 ubiquitinates TFEB downstream of IKK phosphorylation and that this process occurs in normal cell growth conditions, presumably to keep TFEB levels in tight control.

**Fig. 5.**
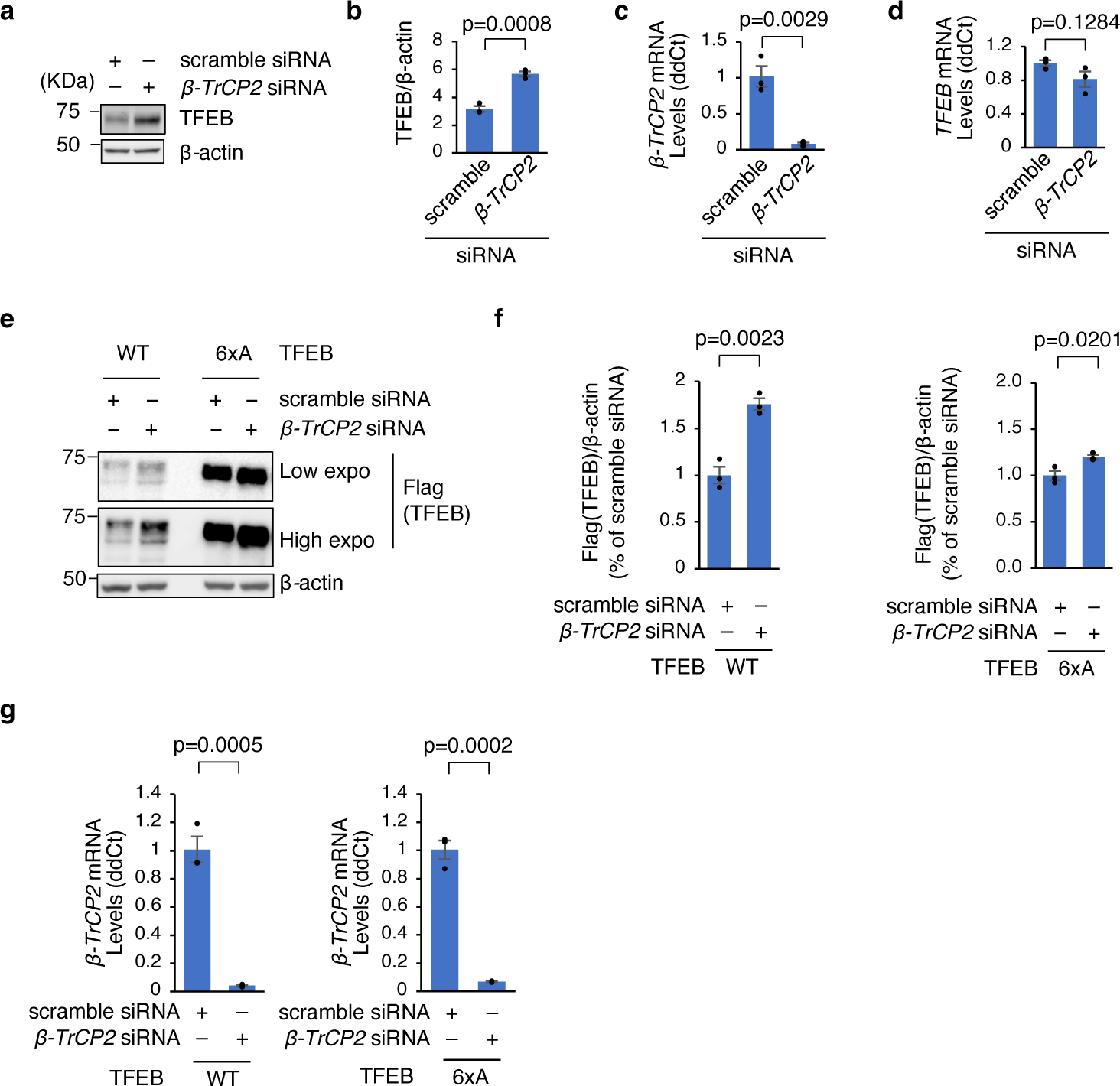
β-TrCP2 promotes TFEB degradation downstream of IKK in normal cell growth conditions. **a** Immunoblot analysis of HEK-293T cells transiently transfected with scramble or *β-Trcp2* siRNA (n = 3 independent experiments) using the indicated antibodies. **b** Quantification of the immunoblots in (**a**), normalized to corresponding β-actin levels. **c** Relative *β-Trcp2* mRNA levels determined by RT-qPCR in samples shown in panel (**a**). **d** Relative *TFEB* mRNA levels determined by RT-qPCR in samples shown in panel (**a**). **e** Immunoblot analysis of HeLa/TFEB-3xFlag and HeLa/TFEB-6xA-3xFlag transiently transfected with scramble or *β- Trcp2* siRNA (n = 3 independent experiments) using the indicated antibodies. **f** Quantification of the immunoblots in (**e**), normalized to corresponding β-actin levels. **g** Relative *β-Trcp2* mRNA levels determined by RT-qPCR in samples shown in panel (**e**). RT-qPCR Results were normalized to the housekeeping gene, *HPRT1*. Results are presented as mean ± S.E.M. *p* values are calculated based on the Student’s *t*-test.

### TFEB ubiquitination and degradation is mediated by phosphodegron-adjacent lysines

β-TrCP proteins interact with their substrates via a phosphodegron that contains two phosphorylated serine residues separated by three or four amino acids [pSxxx(x)pS]^61^. The canonical phosphodegron targeted by β-TrCP has a DpSGxx(x)pS consensus sequence^61^, but β- TrCP also interacts with non-canonical phosphodegrons that have different sets of amino acids around the two phosphoserines^62–64^. Lysine residues that undergo ubiquitination are typically in proximity to their cognate phosphodegrons^65–67^. Although TFEB does not contain the canonical β-TrCP phosphodegron, two of the serines phosphorylated by IKKβ are separated by three amino acids (^423^SPFPS^427^) and are proximal to two consecutive lysines, K430 and K431 (Fig. 6a). To test whether these lysines are the sites of β-TrCP-mediated ubiquitination, we generated HeLa cells stably expressing a TFEB mutant in which both lysines were replaced by arginine residues (HeLa/TFEB-KKRR). Notably, in normal cell growth conditions, the expression levels of the TFEB-KKRR protein were significantly higher than those of WT TFEB (Supplementary Fig. 5a, b), despite no differences between the mRNA levels of the two transgenes (Supplementary Fig. 5c). Thus, similar to the IKK-resistant TFEB-6xA construct, the process leading to increased TFEB-KKRR protein levels must be post-translational. Notably, TFEB-KKRR was insensitive to β-TrCP2 silencing (Fig. 6b-d), thus mimicking the behavior of TFEB-6xA. A ubiquitination assay showed that arginine substitution of lysines 430/431 decreased the levels of ubiquitination of TFEB to levels similar to those observed in WT TFEB upon β-TrCP2 silencing; in addition, silencing of β-TrCP2 did not further modify the ubiquitination levels of TFEB-KKRR, confirming that β-TrCP2-mediated ubiquitination of TFEB requires these phosphodegron- adjacent lysines (Fig. 6e, f). Of note, both TFEB-6xA and TFEB-KKRR mutants exhibited minimal ubiquitination upon proteasome blockage (Supplementary Fig. 6a, b), indicating that the IKK-β-TrCP2-mediated phosphorylation-ubiquitination cascade is the main pathway regulating TFEB proteasomal degradation in normal cell growth conditions. A cycloheximide chase assay demonstrated that mutation of the phosphodegron-adjacent lysines increased TFEB protein stability in normal cell growth conditions (Fig. 6g), again mimicking mutation of the serines within TFEB’s phosphodegron (see Fig. 3d). To assess if K430 and K431 play a role in modulating TFEB stability under stress conditions, we treated HeLa/TFEB and HeLa/TFEB- KKRR cells with increasing doses of LPS for 48 hours. Immunoblot analyses showed that the levels of WT TFEB decreased in a dose-dependent manner following LPS exposure, whereas the levels of TFEB-KKRR showed no significant change (Fig. 6h). These results indicate that LPS-mediated degradation of TFEB requires the phosphodegron-adjacent lysines. To directly connect IKK-mediated phosphorylation of TFEB to modulation of TFEB stability mediated by the phosphodegron-adjacent lysines, we monitored the protein levels of WT TFEB and TFEB- KKRR upon siRNA-mediated silencing of the IKK regulatory subunit IKKγ. Immunoblotting showed that the levels of WT TFEB increased upon IKKγ silencing as expected, whereas the levels of TFEB-KKRR remained unaffected (Fig. 6i, j). Together, these results show that TFEB levels are modulated by a phosphorylation-ubiquitination cascade executed by IKK and β-TrCP2 in normal cell growth conditions as well as upon induced stress.

**Fig. 6.**
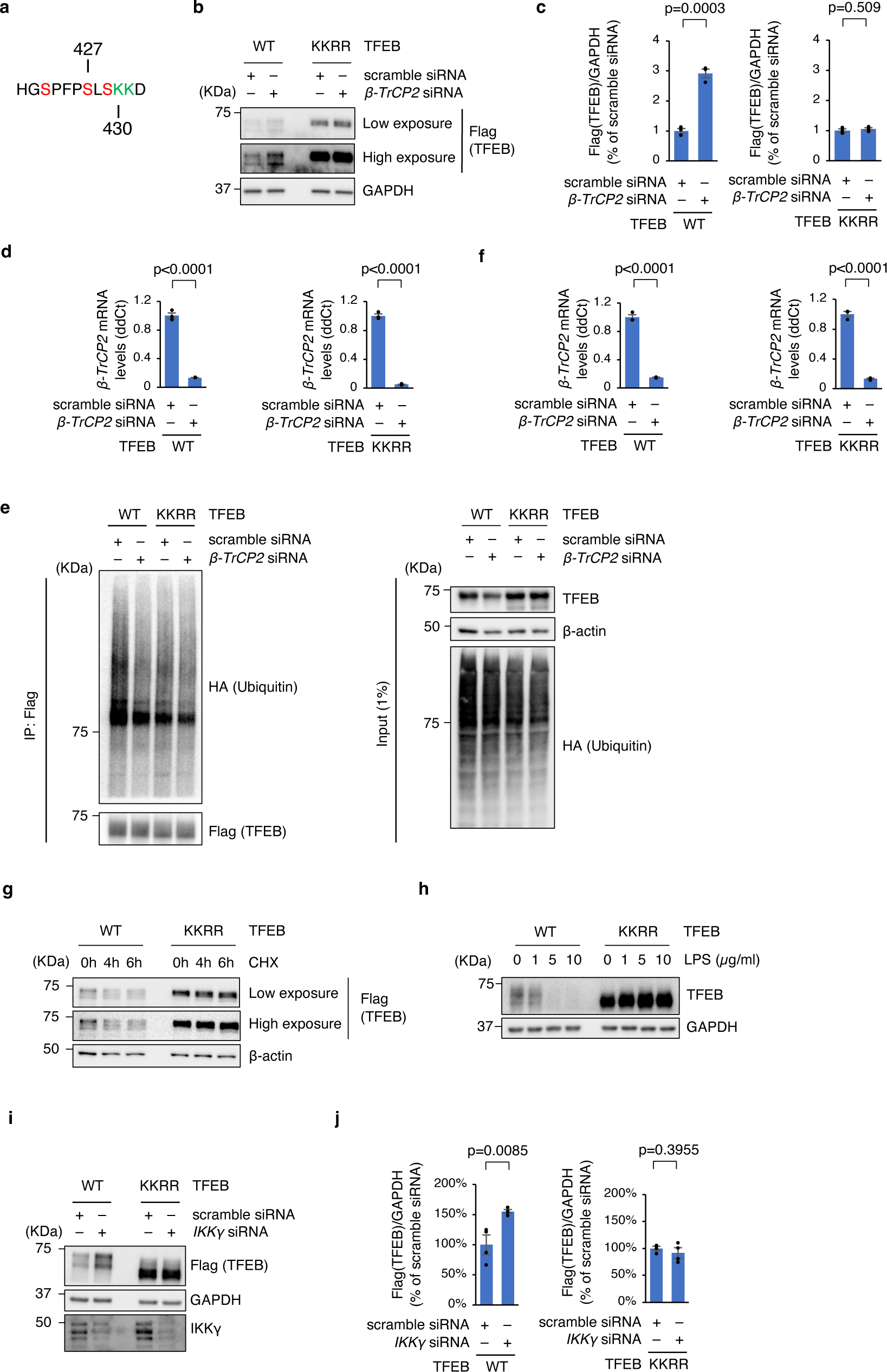
TFEB ubiquitination requires phosphodegron-adjacent lysine residues. **a** Consecutive lysine residues K430 and K431 are adjacent to the TFEB phosphodegron ^423^SPFPSLS^429^. **b** Immunoblot analysis of HeLa/TFEB-3xFlag and HeLa/TFEB-KKRR-3xFlag transiently transfected with scramble or *β-Trcp2* siRNA (n = 3 independent experiments) using the indicated antibodies. **c** Quantification of the immunoblots in (**b**), normalized to corresponding GAPDH levels. **d** Relative *β-Trcp2* mRNA levels determined by RT-qPCR in samples shown in panel (**b**). The expression levels were normalized to the housekeeping gene, *HPRT1*. **e** HEK-293T cells were cotransfected with HA-ubiquitin and TFEB-3xFlag or TFEB- KKRR-3xFlag. 24 h after transfection, the cells were subjected to a second round of transfection with scramble or *β-Trcp2* siRNA. After an additional 48 hrs, TFEB proteins (WT and KKRR) were immunoprecipitated, and ubiquitination levels were assessed using an HA antibody. **f** Relative *β-TrCP2* mRNA levels determined by RT-qPCR in samples shown in panel (**e**). The expression levels were normalized to the housekeeping gene, *HPRT1*. n = 3 technical replicates. **g** HeLa/TFEB-3xFlag and HeLa/TFEB-KKRR-3xFlag were treated with 10 μg/ml cycloheximide (CHX), and the cells were harvested at indicated time points. Immunoblot analyses were performed using the indicated antibodies. **h** HeLa/TFEB-3xFlag and HeLa/TFEB- KKRR-3xFlag were treated with LPS at the specified concentrations for 48h, and the cells were harvested for immunoblot analyses using the indicated antibodies. **i** Immunoblot analysis of HeLa/TFEB-3xFlag and HeLa/TFEB-KKRR-3xFlag transiently transfected with scramble or *IKKγ* siRNA (n = 4 independent experiments) using the indicated antibodies. **j** Quantification of the immunoblots in (**i**), normalized to corresponding GAPDH levels. Results are presented as mean ± S.E.M. *p* values are calculated based on the Student’s *t*-test.

### Inactivation of TFEB degron does not interfere with TFEB nuclear translocation and activation of clearance pathways

Next, we investigated whether degron-dependent modifications of TFEB have a role in TFEB nuclear translocation or TFEB’s ability to modulate clearance pathways. To this aim, we first investigated whether TFEB-6xA and TFEB-KKRR proteins are responsive to starvation and mTORC1 inhibition, stimuli that are both known to promote TFEB nuclear translocation. Confocal microscopy of HeLa/TFEB, HeLa/TFEB-6xA, and HeLa/TFEB-KKRR cells showed that inactivation of either component of TFEB degron did not alter TFEB subcellular distribution or responsiveness to starvation and mTORC1 inhibition (Fig. 7a, b), a result confirmed by immunoblotting after subcellular fractionation (Fig. 7c). Next, we tested the ability of the three constructs to promote the transcription of TFEB target genes. To allow a fair comparison, we did not use the stable clones because they express TFEB at very different levels as shown above. We noticed that transiently expressed WT TFEB, TFEB-6xA, and TFEB-KKRR had similar expression levels (Supplementary Fig. 7a, b), presumably because the very high protein levels associated with transient transfection by themselves overwhelm the processing abilities of IKK/β-TrCP2 or their associated machineries. RT-qPCR experiments performed using RNAs extracted from transiently transfected cells (Fig. 7d) showed that the three constructs enhanced the transcription of TFEB target genes similarly, with TFEB-KKRR showing a slightly enhanced effect (Fig. 7e). We have previously shown that TFEB exogenous expression enhances tau fibril uptake and reduces tau spreading in cell lines and mouse models of Alzheimer’s disease^28, 31^. In order to evaluate the potential of phosphorylation/ubiquitination-resistant TFEB constructs to attenuate tauopathy, we conducted co-transfections of HEK293T cells with a V5-tagged tau- P301L construct along with WT TFEB or either TFEB mutant construct. Quantification of total tau and phospho-tau (PHF1) levels upon comparable expression levels of the three TFEB constructs showed similar overall efficiencies in promoting the degradation of both total tau and phospho-tau, with the two TFEB mutants being slightly more effective at reducing total tau levels (Fig. 7f, g). Taken together, these results indicate that the modifications implicated in TFEB phosphorylation/ubiquitination cascade are functionally distinct from the modifications that control TFEB nuclear translocation, transcriptional capabilities, and clearance activity.

**Fig. 7.**
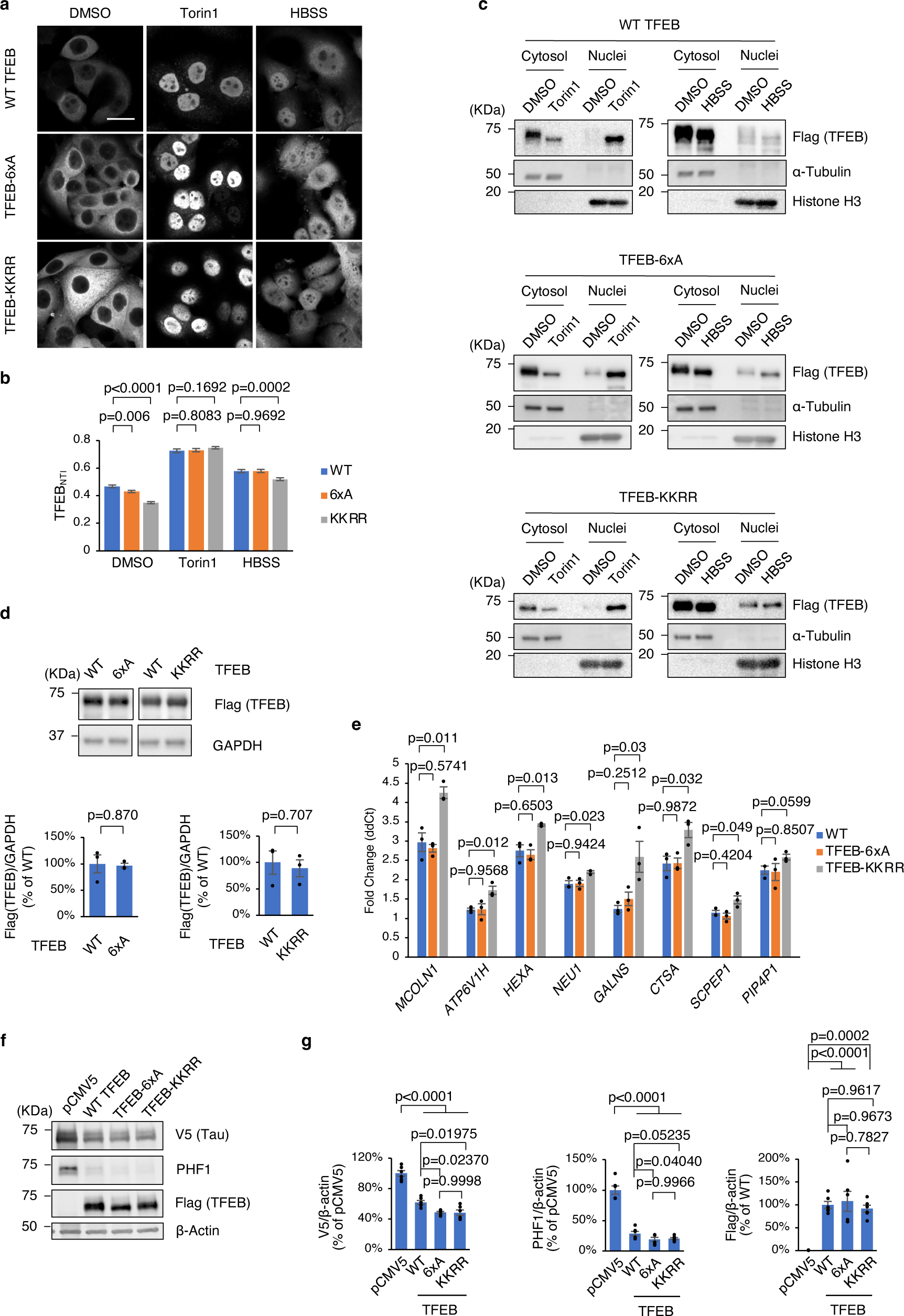
Phosphorylation and ubiquitination of TFEB degron do not interfere with TFEB nuclear translocation and transcriptional capabilities. **a** HeLa/TFEB-3xFlag, HeLa/TFEB- 6xA-3xFlag and HeLa/TFEB-KKRR-3xFlag were treated with DMSO, Torin1 or starved for amino acids (HBSS) for 4 hrs prior to immunofluorescent labelling of TFEB with Flag antibody. Representative images are shown. Scale bar: 30 μm. **b** Quantification of TFEBNTI of cells under the above-mentioned three conditions in (**a**). n > 50 cells per condition. Statistical analysis: Student’s t-test. **c** HeLa/TFEB-3xFlag, HeLa/TFEB-6xA-3xFlag and HeLa/TFEB-KKRR-3xFlag treated as in (**a**) were fractionated into cytosol and nuclei. Immunoblot analyses of each fraction was performed with the indicated antibodies. **d** HEK-293T cells were transfected with 3xFlag tagged WT TFEB, TFEB-6xA or TFEB-KKRR. 24 h after transfection, immunoblot analyses were performed with the indicated antibodies (n = 3 independent experiments) (upper); Quantification of TFEB-6xA and KKRR levels, respectively, relative to WT TFEB levels (lower). Statistical analysis: Student’s t-test. **e** Expression analysis of cells in (**d**). Gene expression was normalized relative to the housekeeping gene, *GAPDH.* Statistical analysis: Student’s t-test. **f** HEK-293T cells were transfected with 3xFlag tagged WT TFEB, TFEB-6xA or TFEB-KKRR, together with V5-tagged full-length human tau with P301L mutation. 48 h after transfection, immunoblot analyses were performed with the indicated antibodies (n = 6 independent replicates for each group). **g** Quantification of total tau, phospho-tau and TFEB levels, respectively. Statistical analysis: one-way ANOVA with Tukey’s multiple comparisons test. Results are presented as mean ± S.E.M.

## DISCUSSION

The lysosome has been historically known for its role as the main catabolic organelle of the eukaryotic cell. A major function of the lysosome is indeed the enzymatic breakdown of macromolecules from all chemical classes (nucleic acids, carbohydrates, lipids, proteins, and their more complex combinations) to dismantle exhausted cellular components and recycle their constituents^68, 69^. Likewise, the lysosome has classically been associated with lysosomal storage disorders (LSDs)—monogenic diseases that arise when any of the lysosomal catabolic pathways is disrupted due to loss-of-function mutations^70–72^. The last decade of research, however, has started to build a solid foundation for a shift in the paradigm that describes the role of the lysosome in cell metabolism. Through the activities of many of its components, the lysosome has gained increased recognition as a central signaling hub that modulates cell metabolism by elaborating responses to information it receives regarding the nutritional status of the cell and its endogenous and exogenous stressors^73^. TFEB is arguably a major player in this hub and the prime effector of lysosome-to-nucleus communication. Not only is TFEB’s subcellular localization directly controlled by mTORC1, the master switch between cellular anabolic and catabolic pathways which itself receives cues from a multitude of nutrient and energy sensors^74^; TFEB also receives direct information from kinases that sit at major signaling nodes of cell metabolism and cell cycle^9–16, 20–23, 75^. Parallel to this paradigm redefinition came the notions that lysosomal homeostasis is central to the pathogenesis of non-LSD late-onset neurodegenerative diseases and that, non-surprisingly, TFEB could be used as a tool to counteract disease progression in degenerative storage diseases by modulating cellular clearance^29, 30, 76, 77^. Exogenous expression of TFEB in animal models of disease has indeed demonstrated TFEB’s ability to ameliorate disease phenotypes relevant to Alzheimer’s disease^26, 28, 31, 78^ and Parkinson’s disease^79–81^ among others. Given that a pharmacologically viable route to enhance the activity of endogenous TFEB has yet to be established, defining the key mechanisms that regulate TFEB levels and function is critically important to lay the foundation for the clinical translation of TFEB-mediated enhancement of cellular clearance.

This work identifies a mechanistically-defined pathway that controls TFEB protein levels by a phosphorylation-ubiquitination-degradation cascade. Our findings reveal that, in normal cell growth conditions, TFEB is subjected to constant proteasomal degradation orchestrated by IKK and β-TrCP2. Phosphorylation of TFEB by IKK generates a phosphodegron that triggers β- TrCP2-mediated degradative ubiquitination of TFEB (Fig. 8). Our data also indicate that TFEB is not a substrate of the β-TrCP2 paralog, β-TrCP1, possibly because the predominant nuclear localization of β-TrCP1^82, 83^ precludes interaction with phosphodegron-bearing TFEB. Remarkably, the mere genetic abolishment of these post-translational events boosts TFEB levels by several fold in the absence of other stimuli or interventions. Also interestingly, the degron targeted by IKK and β-TrCP2 does not overlap with other known signaling modules—a separation that is also reflected functionally, with TFEB phosphodegron mutants still showing full responsiveness to starvation and mTORC1 signaling, ability to transactivate TFEB’s targets genes, and capability to promote tau clearance. As pharmacological inhibitors of IKK are available^84^, these findings provide a novel and unexpected opportunity to modulate TFEB levels for translational studies of lysosomal storage disorders and late-onset neurodegenerative diseases. Future studies on the conditions and stressors that modulate this phosphorylation/ubiquitination cascade could provide additional context on the regulation of TFEB function and its downstream effects and inform on the potential use cases of this pathway in the development of therapeutic avenues. It will also be important to establish the role of the serine residues other than S427 in the phosphodegron. It should be pointed out that the mass-spectrometry data are limited to approximately the signal-to-noise of the MS2 spectra. Thus, under the *in vitro* conditions used here, IKKβ seems to have greater activity toward S427 than S423 and S429 by at least approximately two orders of magnitude. The absence of MS2 signals for S423 or S429 phosphorylation and the absence of MS1 precursors for diphosphorylated forms of these peptides suggests that IKKβ-mediated diphosphorylated-TFEB likely arises from a combination of S427ph with distal sites of phosphorylation. Indeed, our mutational analyses and subsequent experiments have implicated S401, S397 and S399 as additional sites; however, these are located on a very large (∼6 kDa) and acidic (pI ∼4) peptide fragment that proved refractory to proteomic analysis. It is also important to point out that, while generally conserved in vertebrates, TFEB’s phosphodegron does not appear to be present in other members of the microphthalmia/transcription factor E (MiT/TFE) family^85^, which marks an important difference with other described signaling modules of TFEB^12–14, 16, 23^. Whether or not this distinction amongst transcription factors with some functional overlap can be leveraged to design TFEB- targeted therapies warrants future investigation.

**Fig. 8.**
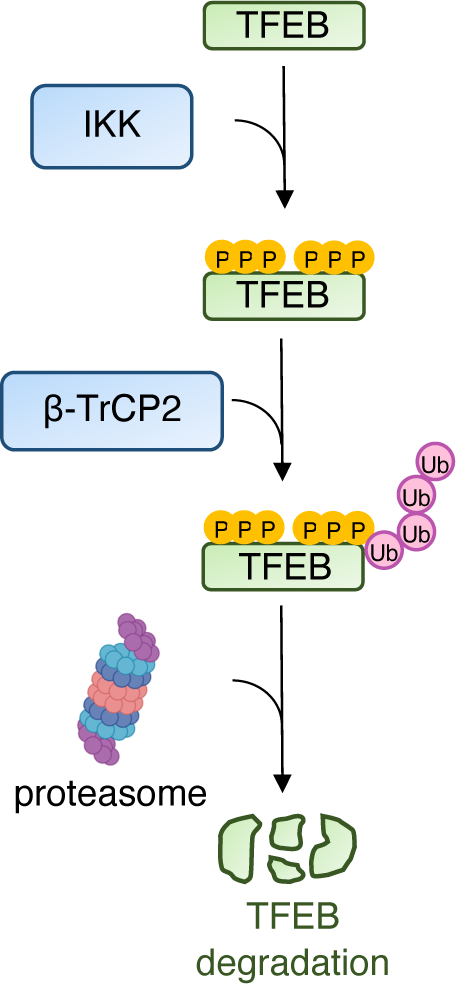
Schematic of the proposed phosphorylation/ubiquitination/degradation mechanism. Phosphorylation of TFEB by IKK triggers ubiquitination executed by β-TrCP2, ultimately resulting in TFEB degradation by the proteasome.

These findings also significantly expand our understanding of the connections of IKK with the autophagy-lysosomal pathway. IKKβ has been reported to regulate the function of mTORC1, an autophagy inhibitor, by two converging routes—direct phosphorylation of mTOR and of mTORC1’s negative regulator TSC1 in response to TNFα, both resulting in mTORC1 activation^32–34^. Given that active mTORC1 phosphorylates TFEB and promotes its cytosolic sequestration, an overall model of IKK regulation of TFEB can be derived where active IKK exerts a double inhibitory effect on TFEB function—cytosolic sequestration via mTORC1 activity, and proteasomal degradation via β-TrCP2-mediated ubiquitination. Our results indeed prove this dual inhibitory effect, as chemical inhibition of IKK simultaneously increased both TFEB protein levels and TFEB nuclear translocation, whereas elimination of IKK-mediated phosphorylation sites only increased TFEB levels while keeping the IKK-resistant protein sensitive to mTORC1 activity. These orthogonal effects of IKK inhibition on nuclear TFEB availability make this route a strong candidate for future testing of TFEB-mediated enhancement of cellular clearance in LSDs and late-onset neurodegenerative diseases.

In summary, this study defines a phosphorylation-ubiquitination cascade responsible for TFEB degradation executed by IKK and β-TrCP2. This mechanism operates independently of the regulation of TFEB nuclear translocation and does not intersect with the control of TFEB’s transcriptional capabilities. This novel pathway adds a critical layer to our understanding of TFEB biology and may have significant therapeutic implications as it represents a pharmacologically actionable entry point to increase the levels of functional TFEB.

## METHODS

### Cell culture, lentivirus infection, and stable cell lines

HeLa cells (ATCC) and HEK293T cells (ATCC) were cultured in DMEM (Cat# 10013CV, Corning™), supplemented with 10% inactivated FBS (Cat# 16140071, Gibco), 2 mM L-glutamine (Cat# 25030081, Life Technologies), and Penicillin/streptomycin (100 U/mL) (Cat# 15140122, Life Technologies). The cells were maintained at 37°C and 5% CO2. HEK293T Cells were plated at a density of 3 × 10^6^ per 60-mm diameter dish 18 h prior to transfection. HEK293T packaging cells were co-transfected with pLenti expression construct, packaging vector psPAX2 (Addgene#12260) and envelope vector pMD2.G (Addgene #12259) using Lipofectamine™ 3000 Transfection Reagent (Cat# L3000001, Thermo Fisher Scientific). The lentiviruses were harvested 72 hr after transfection. Virus particles were purified by centrifugation at 2,000 × g for 10 minutes, followed by filtration through a 0.22 μm pore size syringe filter. For viral transduction, HeLa cells were seeded at 50–60% confluence and treated overnight with the medium containing freshly harvested lentivirus in the presence of 8 μg/mL polybrene (Cat# TR-1003-G, MilliporeSigma). Selection of stable clones was carried out using puromycin (Cat# A1113802, Thermo Fisher Scientific).

### Materials

Reagents used in this study were obtained from the following sources: Antibodies to TFEB (E5P9M) (Cat# 83010S) (1:1000), Flag (D6W5B) (Cat# 14793) (1:1000), β-actin (Cat# 3700S) (1:20,000), HA (C29F4) (Cat# 3724S) (1:1000), α-tubulin (Cat# 2144S) (1:2000), Histone H3 (Cat #9715S) (1:30,000) and IKKγ Kinase (DA10-12) (Cat# 2695T) (1:1000) were from Cell Signaling Technology; antibodies to IKKβ Kinase (Cat# ab124957) (1:5000), IKKα Kinase (Cat# ab32041) (1:5000) and WDR68 (Cat# ab138490) (1:5000) were from Abcam; antibodies to GAPDH (Cat# 60004-1-Ig) (1:150,000) and IκB (Cat# 10268-1-AP) (1:2000) were from Proteintech; antibodies to Flag (produced in mouse) (Cat# F1804) (1:5000), Flag (produced in Rabbit) (Cat# F7425) (1:2000) and c-Myc (Cat# C3956) (1:1000) were from Sigma-Aldrich; antibodies to TrCP1 (C-6) (Cat# sc-390629) (1:1000) and STUB1 (Cat# sc- 133066) (1:1000) were from Santa Cruz; antibodies to V5 (Cat# MCA1360GA) (1:1000) was from Bio-Rad; antibodies to PHF1 antibody (1:1000) was a gift from Dr. Peter Davies (Feinstein Institute for Medical Research). Chemicals: Torin 1 (Cat# 424710) was from Tocris; DMSO (Cat# D8418), Tumor Necrosis Factor-α (TNFα) (Cat# H8916) and cycloheximide (Cat# 01810- 1G) were from Sigma-Aldrich; lipopolysaccharide (LPS) (Cat# tlrl-smlps) was from InvivoGen; carfilzomib (Cat# 17554) was from Cayman Chemical; Protease Inhibitor Cocktail (Cat# P3100- 010) and Phosphotase Inhibitor Cocktail (Cat# P3200-010) were from Gendepot. Opti-MEM (Cat# 31985070) and HBSS (Cat# 14025092) were from Gibco.

### Plasmids and mutagenesis

PCR-amplified human TFEB-GFP was subcloned into pCDH-EF1. PCR-amplified human TFEB-3xFlag was subcloned into a pcDNA3.1 vector. Mutant human TFEB-3xFlag constructs (S3A, S227A, S332A, S334A, S401A, S423A, S441A, S463A, S427A, S429A, S452A, S455A, S462A, S469A, S447A, S459A, S467A, S295A, S389A, S397A, S399A, S466A, S281A, S302A, S352A, 6xA, and KKRR) were generated using the Q5® Site- Directed Mutagenesis Kit (Cat# E0554). TFEB (53-218)-3xFlag, TFEB (219-476)-3xFlag, TFEB-6xA-3xFlag, and plasmids encoding β-TrCP1, DCAF7, STUB1, and myc-β-TrCP2 were synthesized by Twist Bioscience. HA-Ubiquitin (Cat# 17608), Flag-CA-IKKβ (S177E, S181E) (Cat# 11105), Flag-KD-IKKβ (K44M) (Cat# 11104) and HA-β-TrCP2 (Cat# #36969) were from Addgene. Flag-CA-IKKα (S176E, S180E) and Flag-KD-IKKα (K44A) were generated by subcloning PCR-amplified CA/KD-IKKα into a pcDNA3.1 vector. The tau P301L-V5 construct was a gift of Dr. Leonard Petrucelli (Mayo Clinic, Jacksonville, Florida, USA).

### Human kinome screen

All drug treatment, washing, fixing and Hoechst staining steps were performed using the SAMI-integrated automated screening system (Beckman Coulter). HeLa TFEB-GFP cells were seeded at a density of 7,000 cells per well in 96-well plates using a Multidrop and incubated at 37°C in a humidified 5% CO2 incubator for 12 hours prior to treatment. On each plate, four wells were seeded with parental HeLa cells for imaging background controls, eight wells of cells were treated with 0.1% DMSO as drug vehicle controls, and four wells of cells were treated with 250 nM Torin1 as nuclear translocation positive controls. For the remaining wells, each well was treated with one of the 436 drugs within the L1200 library from SelleckChem. Each drug was applied under five different conditions: 10 μM for 6 hours, 3 μM for 6 hours, 1 μM for 6 hours, 10 μM for 1 hour, and 3 μM for 1 hour. After treatment, the cells were fixed with 4% paraformaldehyde at room temperature for 20 minutes and incubated with 2 μM Hoechst 33342 overnight at 4°C. The cells were washed with PBS, and the plates were sealed. Imaging was performed using the IN Cell 2000 Analyzer (GE Healthcare Life Sciences) with FITC (for TFEB-GFP) and DAPI (for nuclei) channels at 20X magnification. Sixteen fields were captured for each well. The acquired images were analyzed with the IN Cell 1000 Workstation analyzer software. DAPI channel images were used to segment the nuclei and identify individual cells for TFEB-GFP intensity measurements. Intensity analysis was conducted by measuring the average intensity of the whole cell. Drugs that yielded intensities greater than the average intensity for all 436 drugs plus four times the standard deviation were considered as outliers and were excluded from further analysis. The Multi Target Analysis Module was used to measure the average nuclear intensity (N) and the average intensity within a 2 µm collar (C) surrounding the nucleus, as the cytoplasmic sampling region.

### Kinome Data analysis

The TFEB nuclear translocation index (TFEBNTI) was calculated as N/(N+C), which ranges from 0 to 1, and was divided into 100 equal intervals. The frequencies of cells falling into each interval were calculated, resulting in 100 data points for each drug, which constitute the distribution curve for each drug and are referred to as the frequency data points. To assess the similarity of the 436 distribution curves to the control distribution curves of DMSO and Torin1, a heatmap was generated using python. The similarity was quantified using the formula: DDMSO * DTorin1, where DDMSO represents the similarity to the DMSO control curves and DTorin1 represents the similarity to the Torin1 control curves. In addition, intensity analysis was conducted by measuring the average intensity of the whole cell. To cluster components based on the frequency data points, several steps were taken to ensure proper normalization and clustering. Initially, min-max normalization was applied to each drug’s frequency data points. This step scaled the values to the range of 0 to 1. Subsequently, we applied quantile normalization to the control components’ data (DMSO and Torin1), which rescaled the frequency data points to a unified scale among all controls and mitigated variations in measurement scales. Following this, frequency data points of all other drugs were adjusted. This adjustment was performed based on the average impact of the quantile normalization on the control data for the associated plates. We then combined the data from different conditions into a single data table for ease of analysis. To identify the relationships between drugs, we computed cosine similarities (distances). These values were utilized in hierarchical clustering, where we implemented single-linkage clustering. In this methodology, the distance between clusters is defined as the shortest distance between any two points in the clusters. In the resulting matrix, each entry signifies the correlation value between the adjusted frequency data points of the two corresponding drugs. Moreover, we calculated the distances between each drug and the mean frequency data points of corresponding Torin1 and DMSO values. These distances were computed using the correlation values that were previously calculated for each drug in relation to all other drugs.

### Transient transfection

Cells were seeded in six-well plates at 70% confluence 12 h prior to transfection. Transfection was performed using Jetprime Transfection Reagent (Cat# 101000027, Polyplus) according to the manufacturer’s protocols. Cells were harvested after 24-48 hours.

### RNA interference and quantitative PCR

HEK293T cells were seeded in six-well plates at 30% confluence 12 h prior to transfection. Cells were transfected with a total of 20 nM siRNAs using the Lipofectamine RNAiMAX transfection reagent (Cat# 13778075, Invitrogen) according to the manufacturer’s protocols. To enhance gene silencing efficiency, two individual siRNAs were used for one gene target. Cells were harvested 48 h or 72 h after transfection. Silencer® Select Negative Control #1 siRNA (Cat# 4390843) and gene specific siRNAs for *β-TrCP2* (Cat# 4392420, assay ID: s23487 and s23485), *IKKβ* (Cat# 4392420, assay ID: s223926 and s7263), *IKKα* (Cat# 4390824, assay ID: s3077 and s3078) and *IKKγ* (Cat# 4390824, assay ID: s16186 and Cat # 4392420, assay ID: s533388) were obtained from Thermo Fisher Scientific. For expression studies, RNA was extracted from cells using the NucleoSpin® RNA kit (Cat# 740955.50, Takara) according to the manufacturer’s instructions. The concentration and quality of the extracted RNAs were assessed using a Nanodrop ONE^c^ (Thermo Scienfic) spectrophotometer. For cDNA synthesis, 1 µg of RNA was used per sample using the QuantiTect Reverse Transcription Kit (Cat# 205311, Qiagen). The quantitative RT-qPCR reactions were carried out using the PowerUp™ SYBR™ Green Master Mix (Cat# A25742, Applied Biosystems) and the QuantStudio-3 real-time PCR systems with the following conditions: 95°C, 2min; (95°C, 15 s; 55°C, 15 s; 72°C, 60 s) x 40. Primer sequences are listed in supplementary Table 2. Relative RNA levels were normalized against an internal control (*GAPDH or HPRT1*) and were calculated as 2^−ΔΔCT^.

### Cell lysis and immunoblotting

Following completion of the treatment, cells were washed once with cold PBS and lysed using either NP-40 lysis buffer (150 mM NaCl, 1% NP-40, 50 mM Tris-Cl pH 8.0) or RIPA buffer (150 mM NaCl, 1% Triton-X100, 50 mM Tris-Cl pH 8.0, 0.5% sodium deoxycholate and 0.1% SDS) supplemented with 1X protease and phosphatase inhibitors. The total lysates were kept at 4°C with agitation for 1 hour, followed by centrifugation at 13,000 rpm for 15 min at 4°C to remove debris. The protein concentration was measured using the Pierce™ BCA Protein Assay Kit (Cat# 23225, Thermo Scientific), with Pierce™ Bovine Serum Albumin Standard, 2 mg/mL (Cat# 23210, Thermo Scientific) as the standard. Protein samples were prepared by adding Laemmli sample buffer (Cat# 1610737EDU, biorad) and incubating at 95°C for 5 min. The samples were then stored at −80°C until use. Denatured samples were separated by SDS-PAGE and subsequently transferred onto PVDF membranes (Cat# 1620177, Biorad). The membranes were incubated in blocking buffer (5%, w/v, dried skimmed milk in Tris-buffered saline, pH 7.4, and 0.1% Tween 20, TBST), followed by overnight incubation with the indicated primary antibodies, appropriately diluted in the blocking buffer. Subsequently, the membranes were washed three times with TBST and incubated with the appropriate secondary antibodies, either Goat anti-mouse HRP-conjugated antibody (Cat# 1705047, Biorad) or Goat anti-rabbit HRP-conjugated antibody (Cat# 1705046, Biorad) or IRDye® 680RD Goat anti-Mouse IgG Secondary Antibody (926-68070, 1:10000, LI-COR Biosciences) or IRDye® 800CW Goat anti- Rabbit IgG Secondary Antibody (926-32211, 1:10000, LI-COR Biosciences), diluted in blocking buffer, for 1 h at room temperature. After washing, the membranes were detected using SuperSignal™ West Dura Extended Duration Substrate reagent (Cat# A38554, ThermoFisher Scientific). Images were taken with ImageQuant LAS 4000 (GE Healthcare) and quantified by Fiji analysis software. Uncropped scans of all blots are provided in **Supplementary Figure 8.**

### Immunoprecipitation

Protein extraction from cultured HEK293T and HeLa cells was performed with NP-40 lysis buffer. After centrifugation (13,000 rpm/15 min, 4 °C) to remove cell debris, the lysates were pre-cleared using dynabeads (Cat# 10003D, Invitrogen) conjugated with normal mouse IgG (Cat# 12-371, Sigma Aldrich). The pre-cleared lysates were then incubated with dynabeads conjugated with the indicated primary antibodies, with rotation at 4°C for 2 hours. Subsequently, the beads were washed four times with 500 μL of RIPA lysis buffer and eluted in 2x Laemmli buffer at 95°C for 5 minutes.

### Immunofluorescence

HeLa cells were seeded at 50–60% confluence on 24-well glass bottom plates (Cat# P24-1.5H-N, Cellvis) and subjected to either transfection or treatment. Cells were fixed with 4% paraformaldehyde at room temperature for 20 minutes, followed by permeabilization in 0.1% Triton X-100 in PBS for 10 min at room temperature with agitation. Next, cell samples were incubated in a blocking buffer consisting of 5% (w/v) Bovine Serum Albumin and 0.1% saposin. Overnight incubation with Flag antibodies (diluted 1:500 in the blocking buffer) was carried out. Subsequently, the cell samples were washed three times with PBS and incubated with the appropriate secondary antibodies: Goat anti-mouse antibody, Alexa Fluor™ 555 (Cat# A-21422, Invitrogen) or Goat anti-mouse antibody, Alexa Fluor™ 488 (Cat# A-11001, Invitrogen), both diluted 1:400 in blocking buffer, for 1 hour at room temperature. Following this, the cell samples were incubated with 300 nM DAPI (Cat# 28718-90-3, Roche) in PBS for 15 min and washed three times with PBS. A Zeiss cell discoverer 7 microscope and accompanying Zen software were used to acquire confocal images.

### *In vitro* kinase assay phos-tag analysis

3xFlag-tagged WT and mutant TFEB were immunoprecipitated from HeLa cell lysate using Flag antibody-conjugated beads. The immunoprecipitated TFEB proteins were dephosphorylated by λ phosphatase (Cat# P9614, Sigma-Aldrich) following the manufacturer’s instructions. Subsequently, the dephosphorylated TFEB samples were subjected to a kinase assay with 300 nM IKKα (Cat# ab102103, abcam), IKKβ (Cat# 81066, Active Motif), or Akt (Cat# 1775-KS-010, R&D systems) in the presence of 100 μM ATP, 2mM MgCl2, 1 mM DTT, 100 μM Na_3_VO_4_, and 500 μM EDTA. The reaction mixtures were incubated for either 60 min or 120 min at 25°C and stopped by adding 2x Laemmli sample buffer followed by boiling for 5 min. Phos-tag analysis was performed according to the manufacturer’s instructions. Subsequently, the samples were separated by 10% SDS–PAGE with 30 μM Phos-binding reagent acrylamide (APExBIO, F4002) and transferred onto PVDF membranes for immunoblotting analyses.

### Sample preparation and LC-MS analysis

Phosphorylated TFEB protein was prepared as described above. To remove MS-incompatible reagents, TFEB was isolated by acetone precipitation. Dry TFEB was reconstituted in 50 mM ammonium bicarbonate buffer pH 7, reduced using dithiothreitol (Pierce, cat. no. 20291), alkylated using iodoacetamide (Sigma- Aldrich, cat. no. A3221), and digested using sequencing grade trypsin (Promega, cat. no. V5113). TFEB tryptic peptides were dried and reconstituted in LC-MS mobile phase A (2% acetonitrile, Thermo Scientific, cat. no.047138.M6), 98% MilliQ purified water, 0.1% formic acid (Pierce, cat. no. 85178) for LC-MS analysis. We used a Thermo Fisher Ultimate RSLC3000 nanoLC system equipped with a micro silica column packed with 20 cm of ReproSil-Pur C18 resin (120Å, C18-AQ,1.9-μmresin; Dr.Maisch, cat. no. r119.aq) and an Orbitrap Fusion Lumos Tribrid Mass Spectrometer. Our acquisition method includes an untargeted acquisition in parallel with a targeted acquisition for the tryptic phosphopeptides at 406AA-430AA and 406AA- 431AA. This approach allows a general analysis of TFEB tryptic peptides and targeted analysis of select candidate phosphopeptides for further fragmentation that ultimately enables the rigorous localization of the phosphorylation sites. The untargeted acquisition settings include Orbitrap as the detector, top 15 data dependent MS2, charge state selection 2-6, 60K and 30K was set as the Orbitrap resolution for MS1 and MS2 respectively, 250-2000 *m/z* scan range, 30% RF lens, 0.8 *m/z* isolation window, stepped HCD at 28%, 30%, 34% collision energy for fragmentation, and profile as acquired data type. The targeted acquisition involves MS2 acquisition, and the settings include a targeted precursor list of *m/z* 881.73 and *m/z* 924.43, Orbitrap as the detector, 30K Orbitrap resolution, scan range is 110-2000 *m/z*, 30% RF lens, stepped HCD fragmentation with 28%, 30%, and 32% collision energy. These fragmentation parameters are optimized to minimize the loss of phosphate for the targeted phosphopeptides. We used our in-house software to process the LC-MS data using UniProt database P19484 for TFEB^86, 87^. The data shows a clear presence of the intact phosphorylated peptide at 406AA- 430AA with *m/z* 881.7311 (-1.99 ppm error, expected mass of 881.732851 *m/z*). The subsequent MS2 data exhibits multiple ion types supporting phosphorylation localization at S427. Both b and y ions that exclude S427 exhibit only unmodified ions, such as y^2^(S429) and y^3^(L428), and all observed b ions. All fragment ions that include S427 exhibit evidence of phosphorylation. Several fragment ions that include S427, such as y^4^(S427), y^5^(P426), y^12^(P419), y^15^(P416), exhibit intact monophosphorylation directly supporting the localization of phosphorylation at S427. We also manually validated the MS2 fragment ions independently to ensure confident phosphorylation site localization.

### Subcellular fractionation

Cells were cultured to 70% confluence in a 6-well plate. Subcellular fractionation was performed using the Subcellular Protein Fractionation Kit for Cultured Cells (Cat# 78840, Thermo Scientific) following the manufacturer’s protocols. Each fraction was subsequently subjected to immunoblotting analyses.

### Data analysis

The data were presented as the mean ± S.E.M. unless otherwise stated. Statistical comparisons between two groups were performed using two-tailed student unpaired t-test; statistical comparisons between multiple groups were performed using one-way analysis of variance (ANOVA) followed by Tukey’s multiple comparison test. A *p* value < 0.05 was considered statistically significant.

## AUTHOR CONTRIBUTIONS

Y.X. and M.S. conceived the project and designed the experiments. M.N.Y. and N.L.Y. designed mass-spectrometry experiments. Y.X. and M.X.G.I designed high throughput screening experiments. Y.X., J.S., W.X., M.N.Y., K.F.P., M.X.G.I., and Q.W. performed experiments and analyzed the data under the supervision of H.Z., N.L.Y. and M.S. A.J. performed bioinformatic analyses. Y.X. and M.S. wrote the manuscript with input from all authors.

## ACKNOWLEDGEMENTS

We thank Mike Prinsen for assistance with running the high throughput screen. This work was supported by NIH grant P01 AG066606 and grants from the Beyond Batten Disease Foundation (to M.S.).

