## Supplementary Figures for "TFEB degradation is regulated by an IKK/β-TrCP2 phosphorylation-ubiquitination cascade"

**Contents:**

Supplementary Figures 1 to 7.

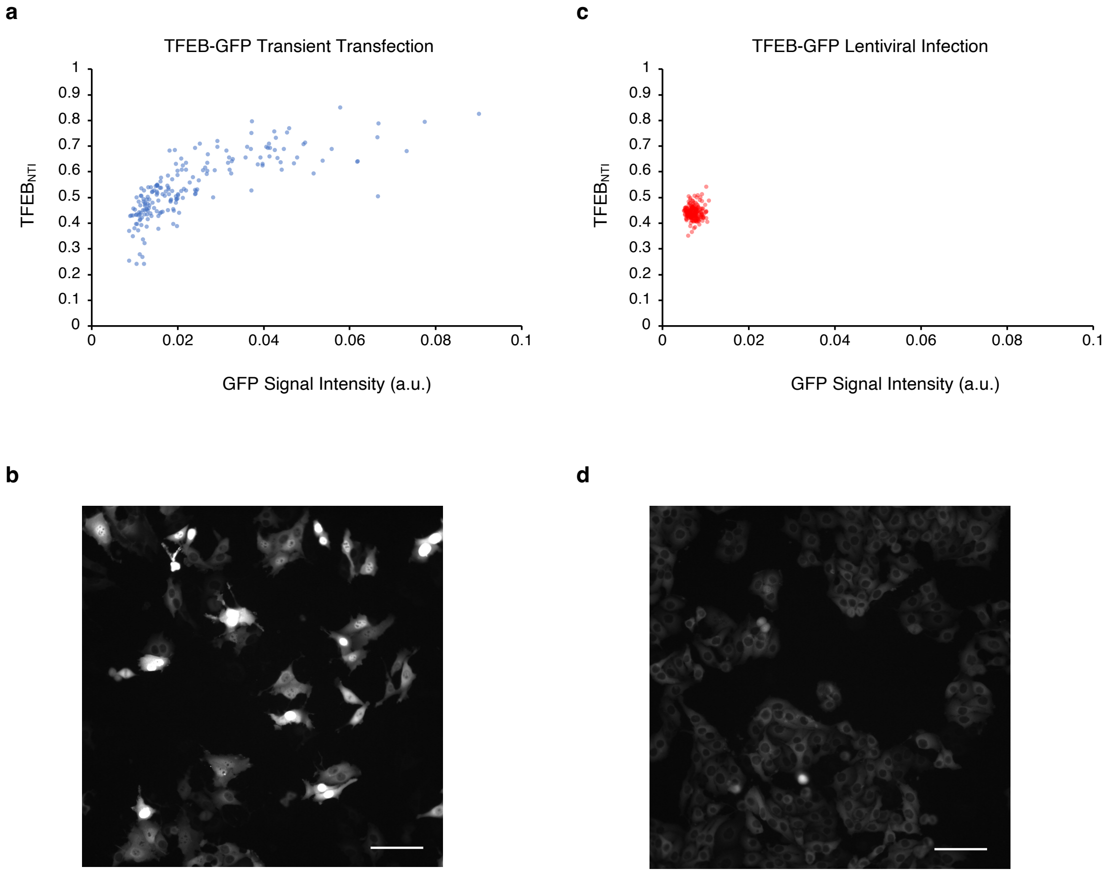


**Supplementary Figure 1**. **Assessment of TFEB nuclear translocation in HeLa cells subjected to TFEB-GFP transient transfection and HeLa/TFEB-GFP**. **a** Dot plot showing the relationship between TFEB-GFP intensity and NT index of individual HeLa cells transiently transfected with TFEB-GFP. **b** A representative image of the analyses in (**a**). **c** Dot plot showing the relationship between TFEB-GFP intensity and NT index of individual HeLa/TFEB-GFP cells. **d** A representative image of the analyses in (**c**). Scale bar in (**b**) and (**d**): 100 μm.

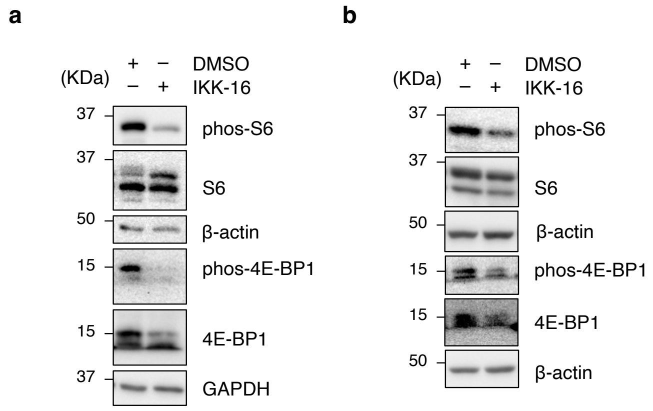


**Supplementary Figure 2**. **Assessment of mTORC1 activity under IKK-16 treatment**. Immunoblot analysis of HEK-293T cells (**a**) and HeLa/TFEB-GFP (**b**) treated with 10 µM IKK-16 or vehicle (0.1% DMSO) for 6 hrs using the indicated antibodies.

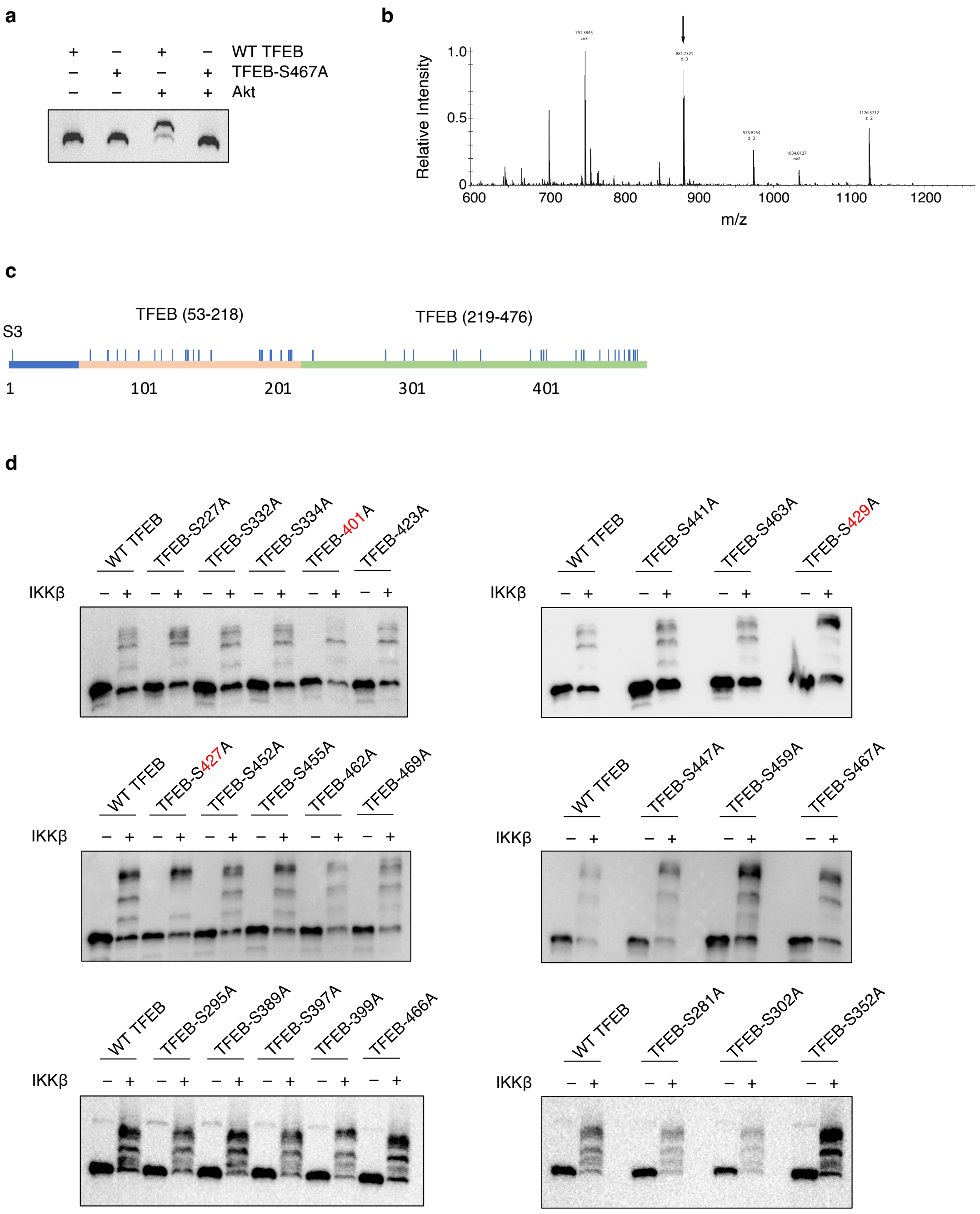


**Supplementary Figure 3**. **Identification of candidate IKK target TFEB serine residues through phos-tag analysis**. **a** *In vitro* kinase assays. Immunoprecipitated TFEB-3xFlag or TFEB S467A-3xFlag were incubated with recombinant Akt and analyzed by phos-tag analysis as described in (**Fig. 2**). **b** Higher energy collisional dissociation (HCD) MS1 mass spectrum of TFEB's tryptic peptide precursor m/z 881.73 corresponding to the amino acid sequence 405AA-

430AA (REDEGPPGYPEPLAPGHGSPFPSLSK). **c** Schematic representation illustrating the TFEB fragmentation strategy. Vertical blue bars represent serine residues. **d** *In vitro* kinase assays. Immunoprecipitated TFEB-3xFlag or 3xFlag-tagged TFEB mutants with specific amino acid substitutions were incubated with recombinant IKKβ and analyzed by phos-tag analysis.

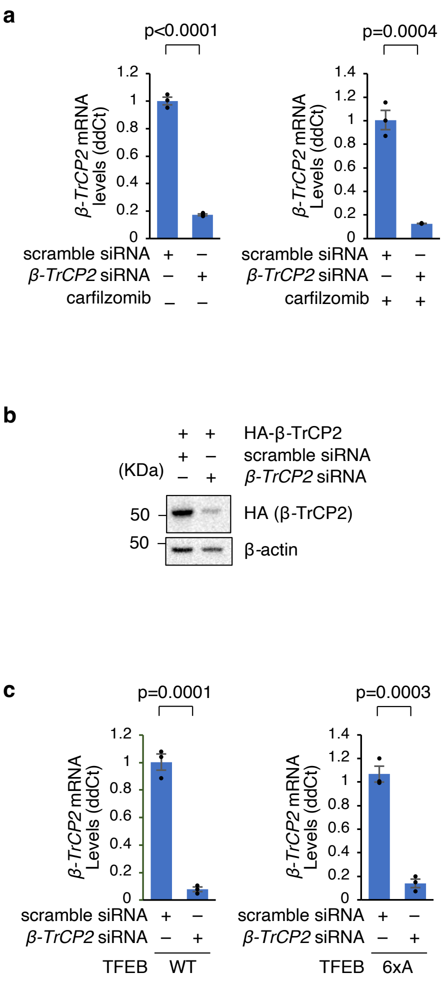


**Supplementary Figure 4**. ***β-TrCP2* siRNA efficiently knocked down HA-tagged β-TrCP2**. **a** Relative *β-Trcp2* mRNA levels determined by RT-qPCR in samples shown in (**Fig. 4b**). **b** HEK-293T cells were transfected with HA-β-TrCP2. 24 h after transfection, the cells were subjected to a second round of transfection with scramble or *β-Trcp2* siRNA. 48 h later, the cells were harvested and β-TrCP2 protein levels were checked using an HA antibody. **c** Relative *β-Trcp2* mRNA levels determined by RT-qPCR in samples shown in (**Fig. 4c**). Results in (**a**) and (**c**) were normalized to the housekeeping gene, *HPRT1* and are presented as mean ± S.E.M. n = 3 technical replicates. *p* values are calculated based on the Student's *t*-test.

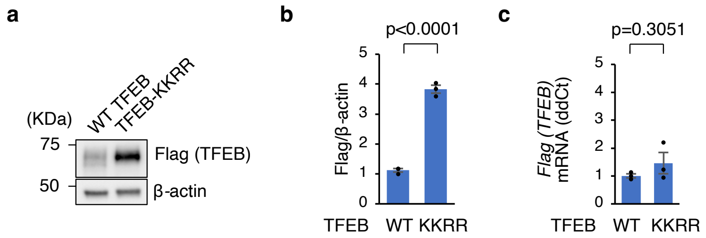


**Supplementary Figure 5**. **Abolishment of β-TrCP2 ubiquitination sites stabilizes TFEB. a** Immunoblot analysis of HeLa/TFEB-3xFlag and HeLa/TFEB-KKRR-3xFlag using the indicated antibodies (n = 3 independent replicates for each group). **b** Quantification of the immunoblots in (**a**), normalized to the corresponding β-actin levels. **c** Relative *TFEB-3xFlag* mRNA levels determined by RT-qPCR in samples shown in panel (**a**). The expression levels were normalized to the housekeeping gene, *GAPDH*. Results in (**b**) and (**c**) are presented as mean ± S.E.M. *p* values are calculated based on the Student's *t*-test.

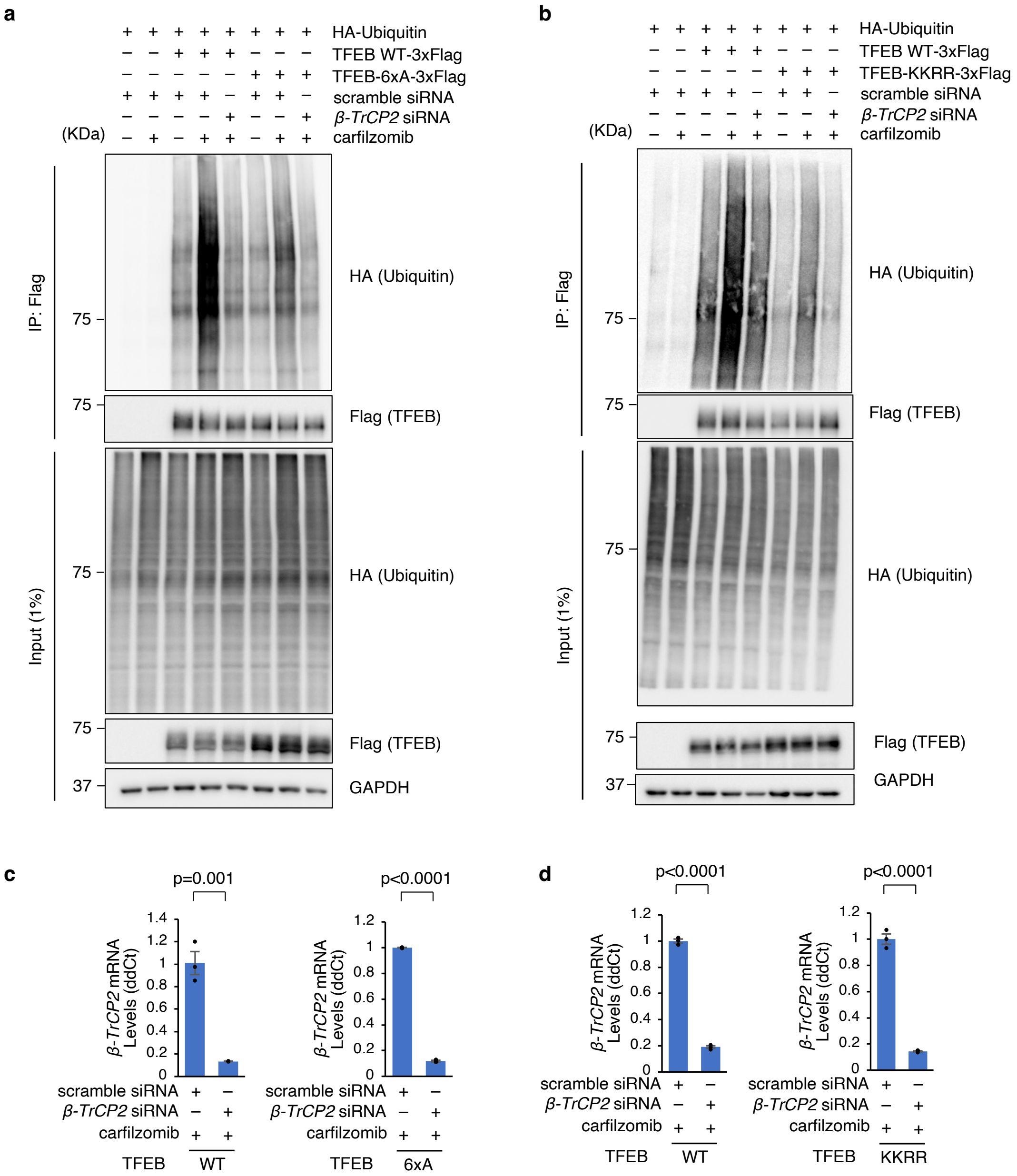


**Supplementary Figure 6**. **Proteasome blockage and β-TrCP2 silencing exert minimal influence on TFEB-6xA and TFEB-KKRR mutants. a, b** HEK-293T cells were cotransfected with HA-ubiquitin and TFEB-3xFlag/ TFEB-6xA-3xFlag (**a**); or cotransfected with HA-ubiquitin and TFEB-3xFlag/ TFEB-KKRR-3xFlag (**b**). 24 h after transfection, the cells were subjected to a second round of transfection with scramble or *β-Trcp2* siRNA. After an additional 28 hrs, the cells were incubated with 100 nM CFZ for 20 h before harvest. TFEB proteins were then immunoprecipitated, and ubiquitination levels were assessed using an HA antibody. **c**

Relative *β-TrCP2* mRNA levels determined by RT-qPCR in samples shown in panel (**a**). **d** Relative *β-TrCP2* mRNA levels determined by RT-qPCR in samples shown in panel (**b**). Results in (**c**) and (**d**) were normalized to the housekeeping gene, *HPRT1* and are presented as mean ± S.E.M. n = 3 technical replicates. *p* values are calculated based on the Student's *t*-test.

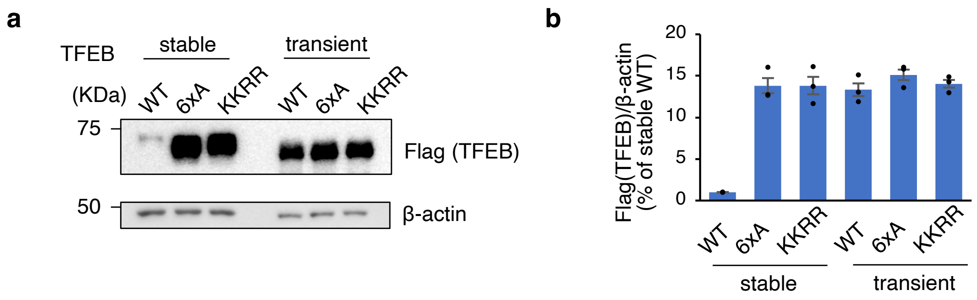


**Supplementary Figure 7**. **Transiently transfected TFEB constructs have comparable expression levels is insensitive to β-TrCP2 or IKKγ silencing. a** Immunoblot analysis of HeLa/TFEB, HeLa/TFEB-6xA, HeLa/TFEB-KKRR, and HeLa cells transiently transfected with WT TFEB, TFEB-6xA, or TFEB-KKRR. **b** Quantification of TFEB expression levels in (**a**) normalized to *β*-actin (n = 3 independent experiments). Data are presented as the mean ± S.E.M. Statistical analysis: Student's *t* test.

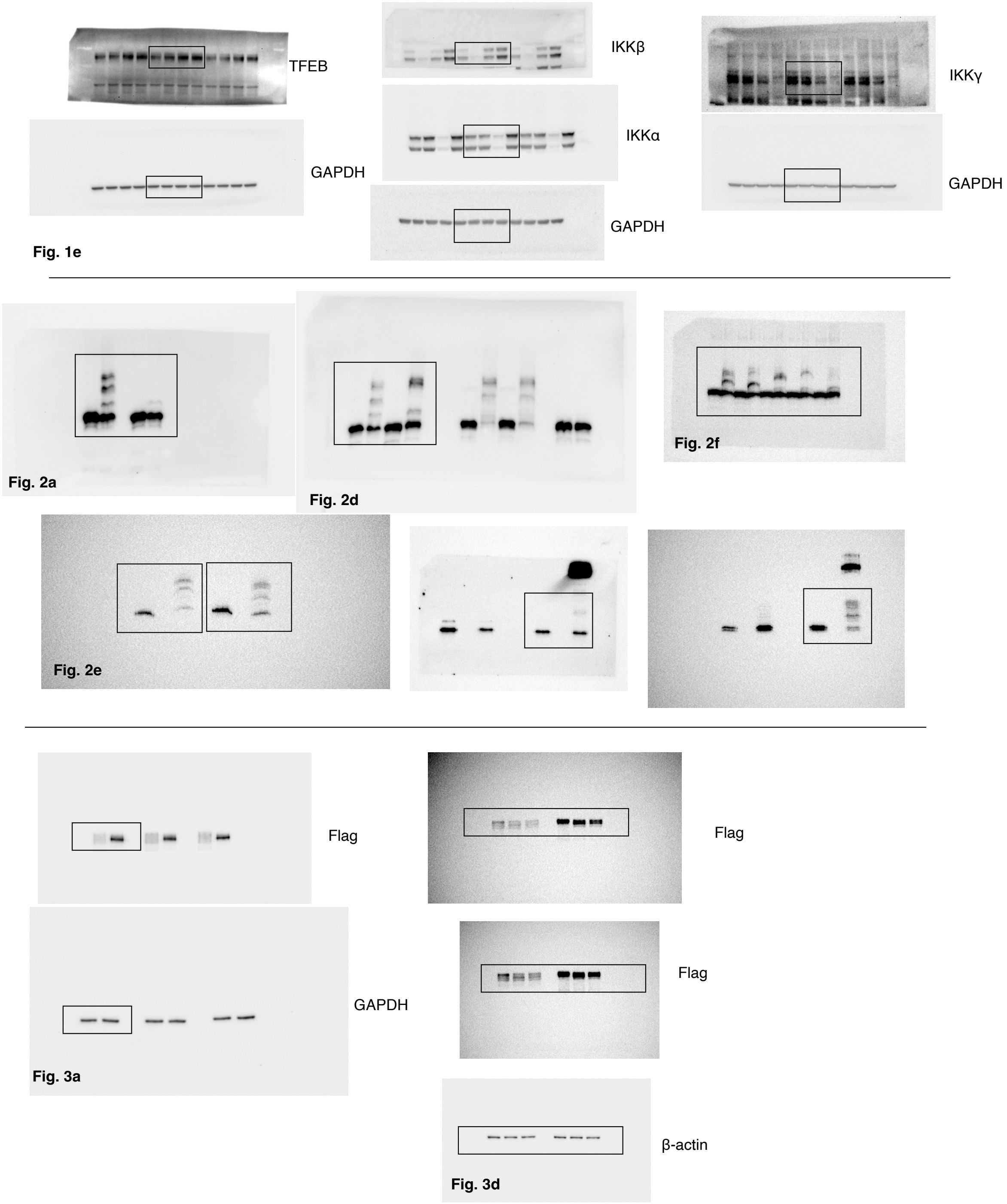


**Supplementary Figure 8**. **Uncropped original western blots.**

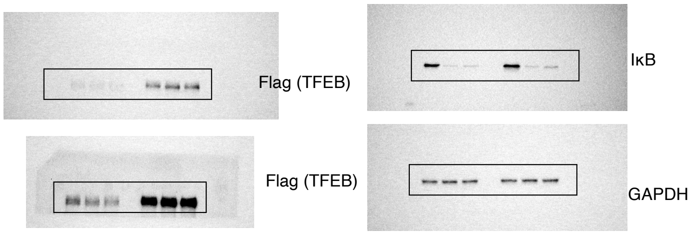

**Fig. 3e**

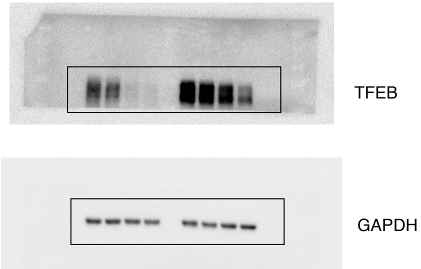

**Fig. 3f**

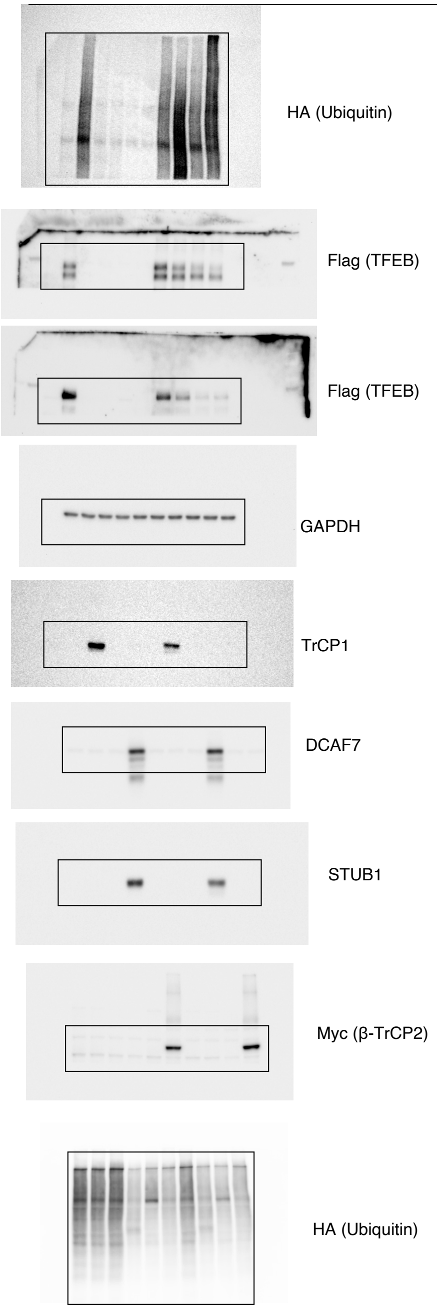

**Fig. 4a**

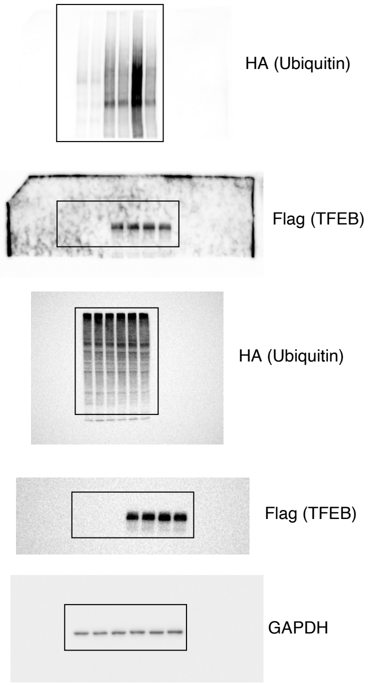

**Fig. 4b**

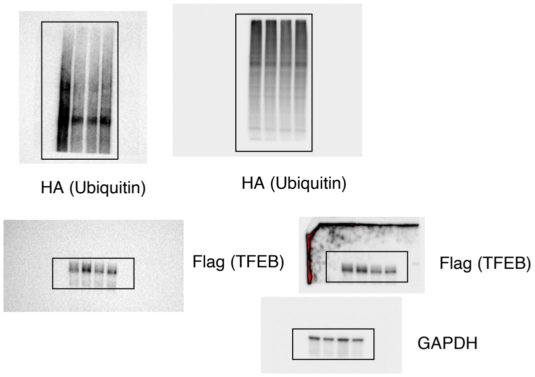

**Fig. 4c**

**Supplementary Figure 8** – ***continued***

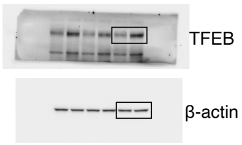


**Fig. 5a**

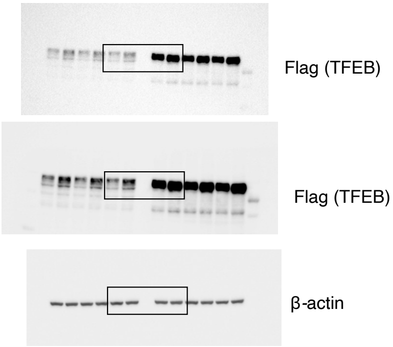


**Fig. 5e**

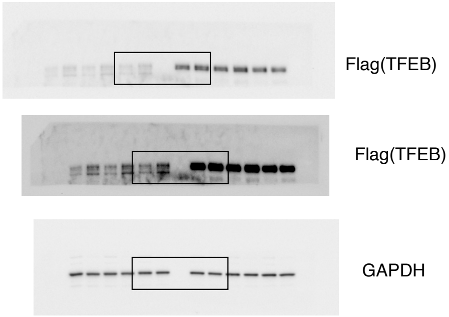


**Fig. 6b**

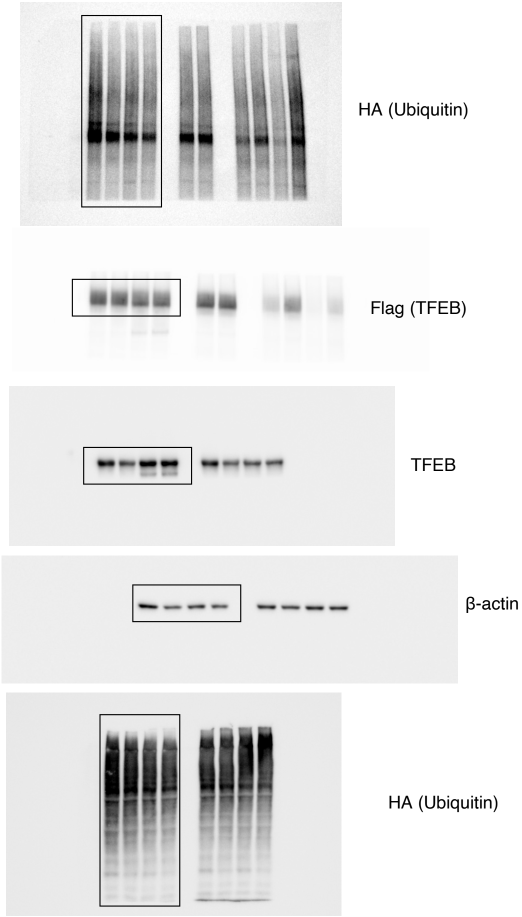


**Fig. 6e**

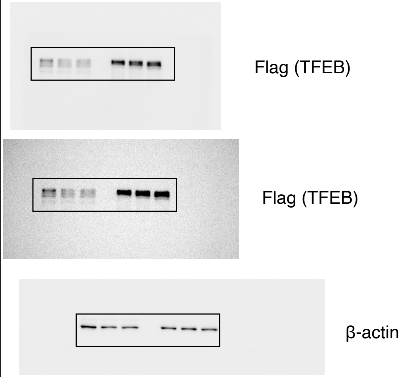


**Fig. 6g**

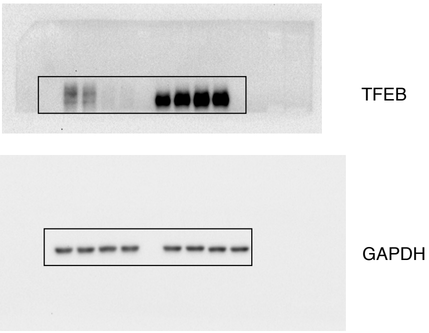


**Fig. 6h**

**Supplementary Figure 8 – *continued***

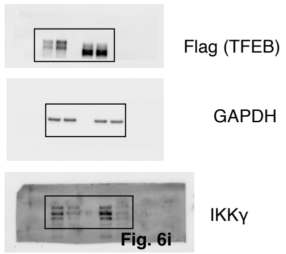


Fig. 6i

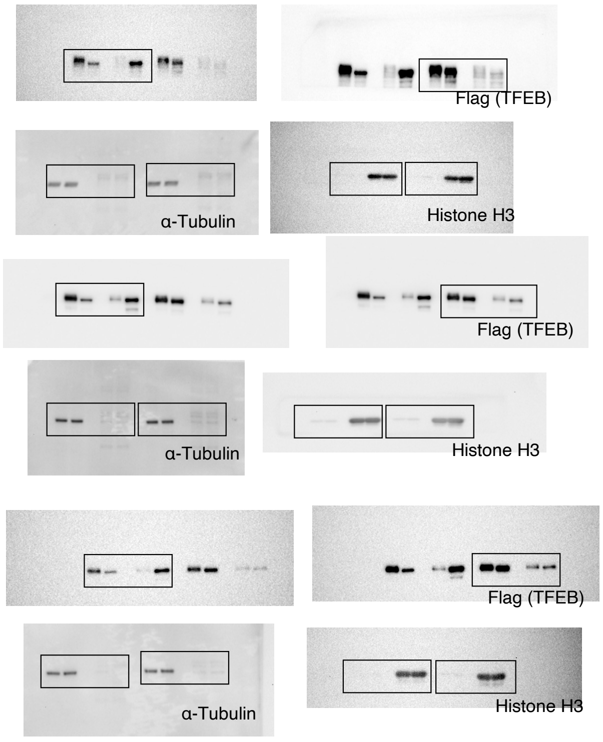


Fig. 7c

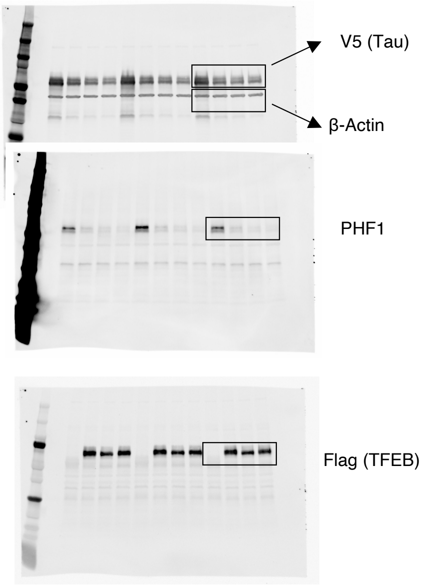


Fig. 7f

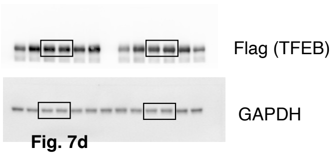


Fig. 7d

**Supplementary Figure 8** – ***continued***

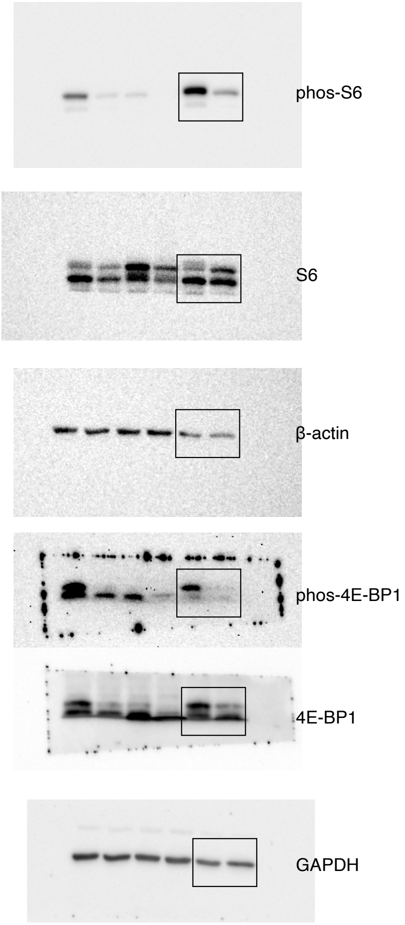

**Supplementary Fig. 2a**

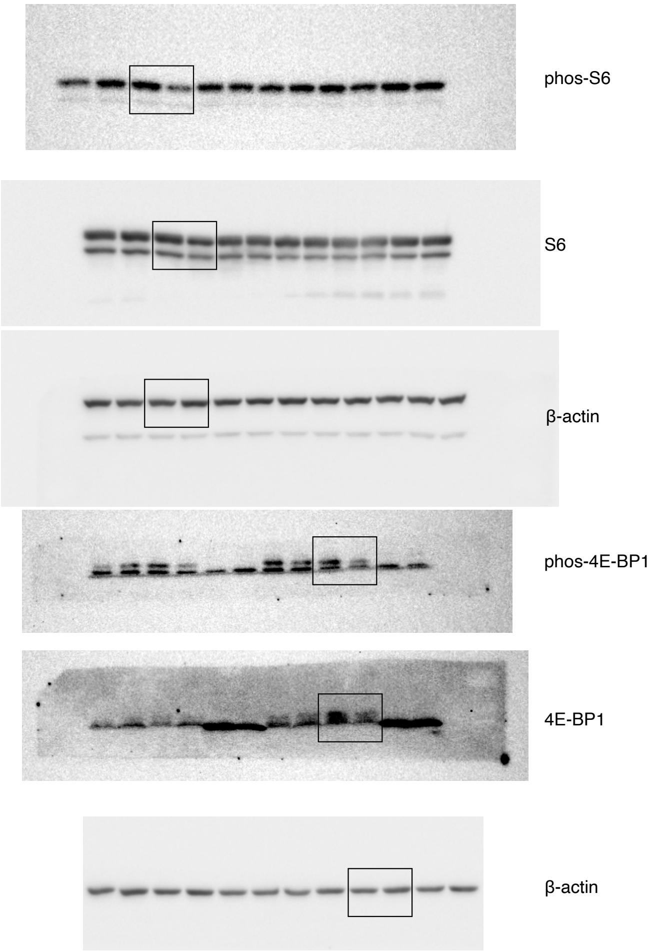

**Supplementary Fig. 2b**

### Supplementary Figure 8 – *continued*

**a**

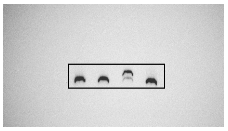

**Supplementary Fig. 3a**

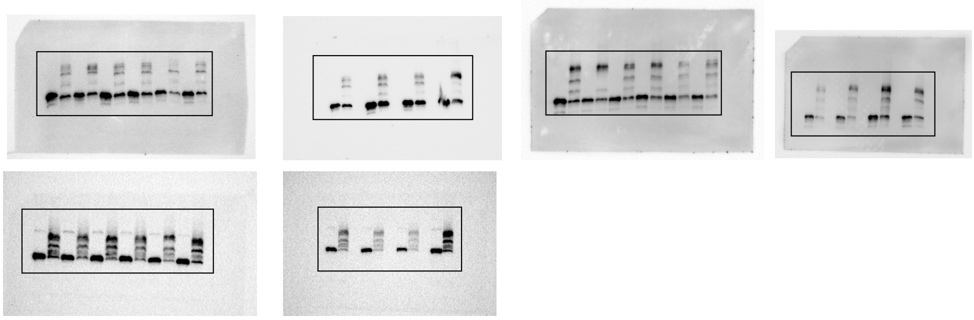

**Supplementary Fig. 3c**

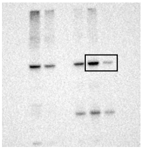

HA (β-TrCP2)

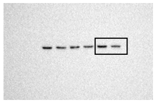

β-actin

**Supplementary Fig. 4**

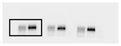

Flag (TFEB)

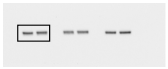

β-actin

**Supplementary Fig. 5**

### Supplementary Figure 8 – *continued*

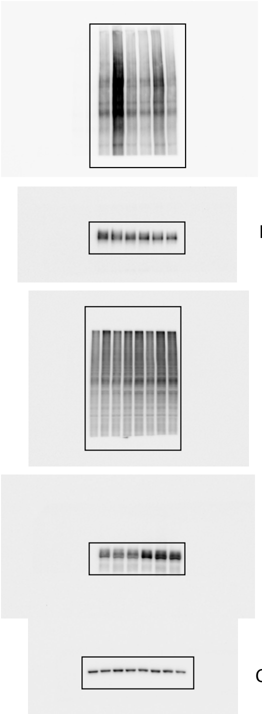

HA (Ubiquitin)

Flag (TFEB)

HA (Ubiquitin)

Flag (TFEB)

GAPDH

**Supplementary Fig. 6a**

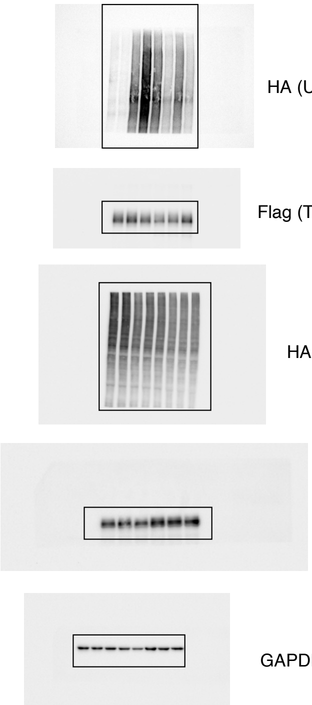

HA (Ubiquitin)

Flag (TFEB)

HA (Ubiquitin)

Flag (TFEB)

GAPDH

**Supplementary Fig. 6b**

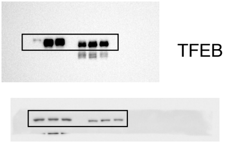

TFEB

β-actin

**Supplementary Fig. 7a**

#### Supplementary Figure 8 - *continued*
