## Supplementary Table 1 for "TFEB degradation is regulated by an IKK/β-TrCP2 phosphorylation-ubiquitination cascade"

**Supplementary Table 1. Distances of kinome screen drugs in hierarchical clustering.**

| Compound | Average Distance from DMSO | Average Distance from Torin1 |
|---|---|---|
| pik-75 | 11.62 | 13.08 |
| pacritinib (sb1518) | 11.44 | 13.06 |
| wye-354 | 12.81 | 11.67 |
| pp121 | 10.92 | 13.65 |
| r428\|bgb324 | 10.96 | 13.57 |
| gdc-0941 | 10.87 | 13.67 |
| ikk-16 (ikk inhibitor vii) | 11.59 | 12.75 |
| mk-8776 (sch 900776) | 10.56 | 13.94 |
| ldk378 | 9.99 | 14.73 |
| ym201636 | 12.71 | 11.58 |
| ly2835219 | 10.64 | 13.73 |
| cudc-907 | 9.59 | 15.14 |
| azd5438 | 11.86 | 12.20 |
| ch5132799 | 15.26 | 9.48 |
| amg-458 | 14.70 | 9.82 |
| everolimus (rad001) | 9.73 | 14.82 |
| nvp-bhg712 | 13.86 | 10.37 |
| osi-027 | 14.58 | 9.80 |
| gsk461364 | 9.48 | 14.96 |
| gsk1059615 | 15.66 | 9.02 |
| zotarolimus(abt-578) | 8.46 | 16.11 |
| bkm120 (nvp-bkm120, buparlisib) | 8.46 | 16.07 |
| imd 0354 | 8.28 | 16.40 |
| staurosporine | 7.99 | 16.85 |
| temsirolimus (cci-779, nsc 683864) | 8.20 | 16.35 |
| pp242 | 16.75 | 7.78 |
| rapamycin (sirolimus) | 7.68 | 16.85 |
| pd173955 | 7.80 | 16.43 |
| sgi-1776 free base | 7.30 | 17.54 |
| ku-0063794 | 17.14 | 7.45 |
| az20 | 17.34 | 7.31 |
| pf-4989216 | 7.23 | 17.51 |
| zstk474 | 7.23 | 17.32 |
| dacomitinib (pf299804, pf299) | 7.03 | 17.82 |
| azd6738 | 7.19 | 17.38 |
| jnk-in-8 | 7.22 | 17.29 |
| tg100713 | 6.86 | 17.81 |
| fingolimod (fty720) HCl | 6.68 | 18.16 |

| | | |
|---|---|---|
| crenolanib (cp-868596) | 6.79 | 17.79 |
| pi-103 | 17.67 | 6.82 |
| cx-6258 HCl | 6.83 | 17.63 |
| azd9291 | 6.72 | 17.78 |
| azd3463 | 6.38 | 18.63 |
| birb 796 (doramapimod) | 6.42 | 18.34 |
| urmc-099 | 6.53 | 18.04 |
| bi 2536 | 6.15 | 18.52 |
| chir-124 | 6.33 | 18.00 |
| cabozantinib (xl184, bms-907351) | 6.04 | 18.79 |
| nu7441 (ku-57788) | 6.01 | 18.81 |
| etp-46464 | 18.73 | 6.04 |
| ponatinib (ap24534) | 5.97 | 18.91 |
| at9283 | 5.94 | 18.66 |
| at7867 | 5.78 | 19.20 |
| crizotinib (pf-02341066) | 5.75 | 19.02 |
| tyrphostin ag 879 | 5.72 | 19.05 |
| degrasyn (wp1130) | 5.66 | 19.23 |
| vs-5584 (sb2343) | 19.02 | 5.68 |
| pha-665752 | 5.79 | 18.50 |
| tak-901 | 5.61 | 19.10 |
| cyc116 | 5.42 | 19.40 |
| bi-847325 | 5.40 | 19.04 |
| azd2014 | 18.96 | 5.41 |
| pik-93 | 5.33 | 19.23 |
| ly294002 | 5.24 | 19.53 |
| milciclib (pha-848125) | 5.25 | 19.42 |
| ve-821 | 5.18 | 19.62 |
| tyrphostin 9 | 5.19 | 19.51 |
| palbociclib (pd-0332991) HCl | 5.12 | 19.65 |
| gsk690693 | 5.12 | 19.60 |
| nu6027 | 5.06 | 19.72 |
| cyt387 | 5.04 | 19.50 |
| pd173074 | 4.96 | 19.67 |
| way-600 | 19.41 | 4.96 |
| tg101348 (sar302503) | 4.88 | 19.69 |
| cp-673451 | 4.85 | 19.70 |
| gdc-0032 | 4.77 | 20.01 |
| ast-1306 | 4.70 | 20.08 |
| aurora a inhibitor i | 4.60 | 20.19 |
| nvp-bsk805 2HCl | 4.52 | 20.45 |

| | | |
|---|---|---|
| foretinib (gsk1363089) | 4.52 | 20.43 |
| ly2090314 | 4.54 | 20.25 |
| sunitinib malate | 4.63 | 19.62 |
| cct137690 | 4.46 | 19.83 |
| wz8040 | 4.28 | 20.64 |
| nilotinib (amn-107) | 4.24 | 20.74 |
| ku-60019 | 4.21 | 20.68 |
| gsk1838705a | 4.13 | 20.89 |
| encorafenib (lgx818) | 4.16 | 20.75 |
| cediranib (azd2171) | 4.11 | 20.80 |
| chir-98014 | 4.04 | 21.10 |
| su6656 | 4.27 | 19.94 |
| sgx-523 | 4.00 | 21.00 |
| xl019 | 4.06 | 20.68 |
| tg101209 | 4.12 | 20.35 |
| chir-99021 (ct99021) | 3.85 | 21.21 |
| g-749 | 3.96 | 20.46 |
| gzd824 | 4.00 | 20.24 |
| nvp-aew541 | 3.80 | 21.01 |
| gsk2334470 | 3.68 | 21.31 |
| jnj-7706621 | 3.77 | 20.74 |
| pha-793887 | 3.78 | 20.66 |
| wp1066 | 3.56 | 21.21 |
| plx-4720 | 3.62 | 20.84 |
| cgk 733 | 3.51 | 21.37 |
| volasertib (bi 6727) | 3.55 | 21.01 |
| a-674563 | 3.52 | 21.09 |
| akti-1/2 | 3.61 | 20.53 |
| afatinib (bibw2992) | 3.44 | 21.19 |
| gne-317 | 21.43 | 3.40 |
| at7519 | 3.46 | 21.03 |
| sns-032 (bms-387032) | 3.35 | 21.63 |
| jnk inhibitor ix | 3.31 | 21.21 |
| chir-99021 (ct99021) HCl | 3.13 | 21.92 |
| nvp-adw742 | 3.13 | 21.78 |
| bms-265246 | 3.11 | 21.72 |
| pf-00562271 | 3.18 | 21.11 |
| pi-3065 | 3.04 | 21.99 |
| r547 | 3.13 | 21.21 |
| gdc-0980 (rg7422) | 21.65 | 3.05 |
| golvatinib (e7050) | 3.00 | 22.04 |

| | | |
|---|---|---|
| ve-822 | 21.30 | 3.09 |
| barasertib (azd1152-hqpa) | 3.09 | 21.28 |
| kw-2449 | 3.03 | 21.70 |
| ddr1-in-1 | 3.08 | 21.27 |
| su11274 | 3.07 | 21.16 |
| hs-173 | 3.05 | 21.16 |
| ldn-214117 | 3.06 | 21.02 |
| su9516 | 3.00 | 21.38 |
| icotinib | 2.93 | 21.67 |
| osu-03012 (ar-12) | 2.90 | 21.81 |
| pf-04691502 | 21.66 | 2.87 |
| mk-2206 2HCl | 2.77 | 22.35 |
| flavopiridol HCl | 2.86 | 21.57 |
| hesperadin | 2.89 | 21.37 |
| del-22379 | 2.76 | 21.99 |
| pd168393 | 2.70 | 22.41 |
| dinaciclib (sch727965) | 2.77 | 21.72 |
| ro3280 | 2.77 | 21.44 |
| pf-431396 | 2.65 | 21.70 |
| bx-912 | 2.58 | 22.11 |
| cnx-2006 | 2.62 | 21.63 |
| ehop-016 | 2.56 | 21.89 |
| quizartinib (ac220) | 2.50 | 22.36 |
| pf-562271 | 2.54 | 21.94 |
| ro-3306 | 2.50 | 22.13 |
| sb590885 | 2.52 | 21.90 |
| pelitinib (ekb-569) | 2.50 | 21.73 |
| go 6983 | 2.43 | 22.35 |
| fiin-2 | 2.49 | 21.72 |
| u0126-etoh | 2.40 | 22.51 |
| motesanib diphosphate (amg-706) | 2.28 | 22.60 |
| gdc-0068 | 2.25 | 22.91 |
| afuresertib (gsk2110183) | 2.24 | 22.87 |
| vatalanib (ptk787) 2HCl | 2.23 | 22.75 |
| enmd-2076 | 2.28 | 22.23 |
| vandetanib (zd6474) | 2.24 | 22.48 |
| pf-477736 | 2.21 | 22.65 |
| p276-00 | 2.19 | 22.64 |
| flavopiridol (alvocidib) | 2.22 | 22.21 |
| gsk650394 | 2.20 | 22.37 |
| bay 11-7082 | 2.18 | 22.35 |

| | | |
|---|---|---|
| byl719 | 2.18 | 22.38 |
| imatinib mesylate (sti571) | 2.15 | 22.58 |
| ipa-3 | 2.08 | 22.75 |
| filgotinib (glpg0634) | 2.05 | 22.69 |
| gne-9605 | 2.04 | 22.66 |
| tak-632 | 2.03 | 22.70 |
| defactinib (vs-6063, pf-04554878) | 2.04 | 22.38 |
| sorafenib | 2.01 | 22.61 |
| gdc-0349 | 22.67 | 2.00 |
| sb239063 | 2.00 | 22.60 |
| gsk621 | 2.00 | 22.48 |
| roscovitine (seliciclib,cyc202) | 1.94 | 23.01 |
| pha-680632 | 1.98 | 22.34 |
| frax597 | 1.90 | 22.77 |
| prt062607 (p505-15, biib057) HCl | 1.86 | 23.25 |
| ku-55933 (atm kinase inhibitor) | 1.87 | 23.13 |
| avl-292 | 1.90 | 22.71 |
| fasudil (ha-1077) HCl | 1.87 | 22.99 |
| osi-906 (linsitinib) | 1.86 | 23.10 |
| h 89 2HCl | 1.86 | 23.20 |
| ssr128129e | 1.88 | 22.83 |
| eht 1864 | 1.88 | 22.79 |
| dorsomorphin (compound c) | 1.90 | 22.52 |
| imatinib (sti571) | 1.82 | 23.36 |
| cgi1746 | 1.81 | 23.25 |
| azd2858 | 1.84 | 22.81 |
| hth-01-015 | 1.84 | 22.67 |
| ink 128 (mln0128) | 22.63 | 1.82 |
| rn486 | 1.77 | 23.36 |
| erk5-in-1 | 1.78 | 23.00 |
| xmd8-92 | 1.77 | 22.97 |
| 6h05 | 1.77 | 22.93 |
| ldc000067 | 1.76 | 22.91 |
| im-12 | 1.75 | 22.84 |
| skepinone-l | 1.71 | 22.85 |
| cct128930 | 1.68 | 23.24 |
| anacardic acid | 1.68 | 22.99 |
| purvalanol a | 1.69 | 22.90 |
| varlitinib | 1.65 | 23.37 |
| gnf-5 | 1.68 | 23.00 |
| gne-0877 | 1.68 | 22.97 |

| | | |
|---|---|---|
| l-685,458 | 1.67 | 22.93 |
| losmapimod (gw856553x) | 1.66 | 22.97 |
| ulixertinib (bvd-523, vrt752271) | 1.67 | 22.64 |
| kn-62 | 1.63 | 23.15 |
| pha-767491 | 1.64 | 23.02 |
| dasa-58 | 1.63 | 23.05 |
| lfm-a13 | 1.63 | 23.05 |
| masitinib (ab1010) | 1.61 | 23.33 |
| wh-4-023 | 1.62 | 23.01 |
| azd2932 | 1.60 | 23.18 |
| lapatinib | 1.56 | 23.43 |
| gf109203x | 1.61 | 22.65 |
| azd4547 | 1.55 | 23.44 |
| wortmannin | 1.54 | 23.47 |
| sns-314 mesylate | 1.55 | 23.20 |
| k-ras(g12c) inhibitor 9 | 1.57 | 22.93 |
| co-1686 (avl-301) | 1.56 | 22.99 |
| azd1208 | 1.54 | 23.37 |
| bikinin | 1.55 | 23.11 |
| kn-93 phosphate | 1.54 | 23.12 |
| linifanib (abt-869) | 1.53 | 23.36 |
| ono-4059 | 1.55 | 23.06 |
| gne-7915 | 1.54 | 23.17 |
| thiazovivin | 1.53 | 23.24 |
| alisertib (mln8237) | 1.55 | 22.87 |
| rigosertib (on-01910) | 1.54 | 22.96 |
| zcl278 | 1.53 | 23.15 |
| entrectinib (rxdx-101) | 1.56 | 22.56 |
| ar-a014418 | 1.53 | 23.02 |
| tg003 | 1.52 | 23.08 |
| azd5363 | 1.47 | 23.64 |
| krn 633 | 1.48 | 23.34 |
| ana-12 | 1.49 | 23.16 |
| azd1080 | 1.47 | 23.40 |
| dovitinib (tki-258) dilactic acid | 1.48 | 23.06 |
| isrib (trans-isomer) | 1.49 | 22.91 |
| ly2784544 | 1.47 | 23.11 |
| 1-azakenpaullone | 1.44 | 23.41 |
| dovitinib (tki-258, chir-258) | 1.45 | 23.24 |
| vx-680 (tozasertib, mk-0457) | 1.46 | 22.99 |
| tie2 kinase inhibitor | 1.44 | 23.42 |

| | | |
|---|---|---|
| gsk2578215a | 1.45 | 23.14 |
| raf265 (chir-265) | 1.44 | 23.33 |
| at13148 | 1.44 | 22.89 |
| bix 02188 | 1.40 | 23.46 |
| blz945 | 1.41 | 23.19 |
| pexmetinib (arry-614) | 1.42 | 22.95 |
| pf-543 | 1.39 | 23.15 |
| bgt226 (nvp-bgt226) | 24.41 | 1.31 |
| 10058-f4 | 1.37 | 23.22 |
| cobimetinib (gdc-0973, rg7420) | 1.35 | 23.51 |
| amg319 | 1.37 | 23.17 |
| bms-345541 | 1.35 | 23.43 |
| wz4003 | 1.35 | 23.29 |
| pazopanib | 1.34 | 23.14 |
| pfk15 | 1.34 | 22.88 |
| bx-795 | 1.33 | 23.09 |
| gsk429286a | 1.29 | 23.46 |
| kx2-391 | 1.29 | 23.36 |
| nintedanib (bibf 1120) | 1.31 | 22.97 |
| bio | 1.30 | 23.05 |
| cnx-774 | 1.29 | 23.10 |
| ly2603618 | 1.26 | 23.65 |
| bosutinib (ski-606) | 1.30 | 22.90 |
| mln8054 | 1.24 | 23.58 |
| axitinib | 1.26 | 23.11 |
| aee788 (nvp-aee788) | 1.25 | 23.21 |
| zm 336372 | 1.23 | 23.63 |
| pf-573228 | 1.23 | 23.20 |
| tivantinib (arq 197) | 1.24 | 22.91 |
| wz3146 | 1.21 | 23.36 |
| bs-181 HCl | 1.20 | 23.47 |
| ac480 (bms-599626) | 1.19 | 23.77 |
| bay 11-7085 | 1.20 | 23.44 |
| osi-930 | 1.18 | 23.67 |
| nvp-bvu972 | 1.19 | 23.52 |
| sb203580 | 1.17 | 23.43 |
| d 4476 | 1.17 | 23.35 |
| ski ii | 1.17 | 23.28 |
| bi-d1870 | 1.17 | 23.14 |
| amuvatinib (mp-470) | 1.15 | 23.52 |
| cep-33779 | 1.15 | 23.71 |

| | | |
|---|---|---|
| genistein | 1.15 | 23.51 |
| ly2228820 | 1.12 | 23.80 |
| sorafenib tosylate | 1.13 | 23.49 |
| dasatinib | 1.11 | 23.71 |
| gsk2636771 | 1.12 | 23.32 |
| pik-293 | 1.11 | 23.49 |
| brivanib (bms-540215) | 1.10 | 23.65 |
| pd0325901 | 1.08 | 23.39 |
| az 960 | 1.10 | 23.01 |
| pht-427 | 1.08 | 23.58 |
| ki8751 | 1.07 | 23.52 |
| danusertib (pha-739358) | 1.10 | 22.99 |
| sotrastaurin | 1.07 | 23.59 |
| telatinib | 1.04 | 23.84 |
| tae226 (nvp-tae226) | 1.05 | 23.51 |
| mubritinib (tak 165) | 1.05 | 23.63 |
| tgx-221 | 1.04 | 23.65 |
| sb202190 (fhpi) | 1.04 | 23.62 |
| pf-4708671 | 1.02 | 23.92 |
| semaxanib (su5416) | 1.03 | 23.33 |
| azd8186 | 1.00 | 24.09 |
| tic10 | 1.01 | 23.73 |
| tws119 | 1.02 | 23.52 |
| amg-900 | 1.03 | 23.25 |
| gdc-0994 | 1.02 | 23.34 |
| lapatinib (gw-572016) ditosylate | 1.01 | 23.71 |
| butein | 0.99 | 23.88 |
| tepotinib (emd 1214063) | 0.99 | 23.83 |
| gsk1904529a | 0.99 | 23.74 |
| refametinib (rdea119, bay 86-9766) | 1.00 | 23.50 |
| pq 401 | 0.99 | 23.72 |
| ipi-145 (ink1197) | 0.98 | 23.46 |
| y-27632 2HCl | 0.97 | 23.70 |
| gnf-2 | 0.97 | 23.65 |
| regorafenib (bay 73-4506) | 0.97 | 23.64 |
| sar245409 (xl765) | 0.97 | 23.43 |
| zm 39923 HCl | 0.96 | 23.74 |
| ibrutinib (pci-32765) | 0.96 | 23.70 |
| pp2 | 0.96 | 23.69 |
| tak-733 | 0.97 | 23.48 |
| hmn-214 | 0.97 | 23.23 |

| | | |
|---|---|---|
| tivozanib (av-951) | 0.95 | 23.70 |
| gefitinib (zd1839) | 0.95 | 23.65 |
| bix 02189 | 0.93 | 23.90 |
| mk-8745 | 0.95 | 23.30 |
| tpca-1 | 0.94 | 23.49 |
| bms-754807 | 0.95 | 23.13 |
| dcc-2036 (rebastinib) | 0.93 | 23.51 |
| bms-777607 | 0.91 | 24.01 |
| piceatannol | 0.91 | 23.80 |
| azd8330 | 0.91 | 23.85 |
| vemurafenib (plx4032, rg7204) | 0.90 | 23.89 |
| erlotinib HCl (osi-744) | 0.91 | 23.75 |
| r406 (free base) | 0.92 | 23.24 |
| vx-702 | 0.89 | 23.86 |
| trametinib (gsk1120212) | 0.91 | 23.45 |
| enzastaurin (ly317615) | 0.89 | 23.90 |
| xl147 | 0.89 | 23.66 |
| apatinib | 0.87 | 24.02 |
| tak-285 | 0.87 | 23.91 |
| cabozantinib malate (xl184) | 0.87 | 23.84 |
| saracatinib (azd0530) | 0.88 | 23.52 |
| a-769662 | 0.83 | 24.06 |
| pik-294 | 0.84 | 23.76 |
| gw441756 | 0.84 | 23.87 |
| wz4002 | 0.82 | 23.85 |
| gdc-0623 | 0.83 | 23.58 |
| gsk2126458 (gsk458) | 23.75 | 0.82 |
| gw5074 | 0.82 | 23.61 |
| acadesine | 0.80 | 24.03 |
| tg100-115 | 0.80 | 23.76 |
| mk-2461 | 0.79 | 23.87 |
| pf-3758309 | 0.81 | 23.15 |
| pf-04217903 | 0.80 | 23.53 |
| cp-724714 | 0.78 | 23.97 |
| wye-125132 (wye-132) | 23.58 | 0.79 |
| vx-745 | 0.78 | 23.80 |
| cudc-101 | 0.78 | 23.51 |
| azd7762 | 0.79 | 23.15 |
| indirubin | 0.77 | 23.80 |
| a66 | 0.76 | 23.98 |
| phenformin HCl | 0.76 | 23.92 |

| | | |
|---|---|---|
| tofacitinib (cp-690550) citrate | 0.75 | 23.73 |
| ap26113 | 0.77 | 23.17 |
| ruxolitinib (incb018424) | 0.74 | 23.70 |
| ph-797804 | 0.73 | 24.02 |
| bms-794833 | 0.73 | 23.92 |
| zm 447439 | 0.74 | 23.66 |
| selumetinib (azd6244) | 0.72 | 23.95 |
| lenvatinib (e7080) | 0.72 | 23.87 |
| tsu-68 (su6668, orantinib) | 0.71 | 24.02 |
| azd6482 | 0.71 | 23.92 |
| ag-1024 | 0.71 | 23.75 |
| pd184352 (ci-1040) | 0.70 | 23.68 |
| pazopanib hcl (gw786034 HCl) | 0.70 | 23.92 |
| rki-1447 | 0.71 | 23.54 |
| cep-32496 | 0.70 | 23.63 |
| mek162 (arry-162, arry-438162) | 0.69 | 23.94 |
| dabrafenib (gsk2118436) | 0.69 | 23.82 |
| bms-536924 | 0.69 | 23.51 |
| asiatic acid | 0.67 | 24.09 |
| brivanib alaninate (bms-582664) | 0.67 | 23.99 |
| ag-18 | 0.67 | 23.96 |
| ag-490 (tyrphostin b42) | 0.66 | 23.90 |
| ro5126766 (ch5126766) | 0.65 | 23.95 |
| 3-methyladenine | 0.66 | 23.71 |
| mk-5108 (vx-689) | 0.65 | 23.65 |
| pp1 | 0.65 | 23.70 |
| mgcd-265 | 0.64 | 23.92 |
| tdzd-8 | 0.63 | 23.84 |
| tofacitinib (cp-690550, tasocitinib) | 0.63 | 23.84 |
| pd98059 | 0.62 | 23.92 |
| r406 | 0.64 | 23.39 |
| whi-p154 | 0.62 | 23.84 |
| cct129202 | 0.62 | 23.63 |
| s-ruxolitinib (incb018424) | 0.61 | 23.78 |
| fostamatinib (r788) | 0.62 | 23.53 |
| azd8931 (sapitinib) | 0.60 | 24.03 |
| as-252424 | 0.60 | 23.93 |
| zm 323881 HCl | 0.61 | 23.86 |
| triciribine | 0.60 | 23.88 |
| gdc-0879 | 0.60 | 23.82 |
| czc24832 | 0.59 | 23.76 |

| | | |
|---|---|---|
| tak-715 | 0.59 | 23.70 |
| pimasertib (as-703026) | 0.58 | 24.02 |
| jnj-38877605 | 0.58 | 24.00 |
| sl-327 | 0.58 | 23.88 |
| sp600125 | 0.57 | 23.87 |
| sb415286 | 0.57 | 23.89 |
| chrysophanic acid | 0.56 | 24.07 |
| osi-420 | 0.57 | 23.80 |
| ag-1478 (tyrphostin ag-1478) | 0.57 | 23.86 |
| fr 180204 | 0.56 | 23.93 |
| as-604850 | 0.55 | 23.91 |
| honokiol | 0.55 | 24.01 |
| zm 306416 | 0.55 | 23.87 |
| sc-514 | 0.55 | 23.76 |
| torin 2 | 24.10 | 0.52 |
| azd8055 | 23.45 | 0.53 |
| az 628 | 0.52 | 23.69 |
| quercetin | 0.50 | 23.82 |
| tcs 359 | 0.46 | 23.76 |
| pd318088 | 0.43 | 24.00 |
| nsc 23766 | 0.43 | 24.01 |
| smi-4a | 0.43 | 23.87 |
| azd1480 | 0.42 | 23.70 |
| tyrphostin ag 1296 | 0.42 | 23.77 |
| anisomycin | 0.42 | 23.58 |
| sb216763 | 0.41 | 23.69 |
| sc79 | 0.37 | 23.97 |
| cal-101 (idelalisib, gs-1101) | 0.36 | 23.82 |
| naringin | 0.33 | 23.78 |
