## Supplementary Table 2 for "TFEB degradation is regulated by an IKK/β-TrCP2 phosphorylation-ubiquitination cascade"

**Supplementary Table 2. List of oligos used in RT-qPCR experiments.**

| Gene name | Forward primer | Reverse primer |
|---|---|---|
| *ATPV1H* | GGAAGTGTCAGATGATCCCCA | CCGTTTGCCTCGTGGATAAT |
| *CTSA* | CAGGCTTTGGTCTTCTCTCCA | TCACGCATTCCAGGTCTTTG |
| *FBXW11* | ACAAGTACATCGTGTCTGCC | TTGTGCCCATTGAGAGTACG |
| *Flag* | AGCTTCAGCATGGAGGA | CACCGTCATGGTCTTTGTAG |
| *GALNS* | TTGTCGGCAAGTGGCATCT | CCAAACCACTCATCAAATCCG |
| *GAPDH* | TGCACCACCAACTGCTTAGC | GGCATGGACTGTGGTCATGAG |
| *HEXA* | CAACCAACACATTCTTCTCCA | CGCTATCGTGACCTGCTTTT |
| *HPRT1* | TGACACTGGCAAAACAATGCA | GGTCCTTTTCACCAGCAAGCT |
| *MCOLN1* | TTGCTCTCTGCCAGCGGTACTA | GCAGTCAGTAACCACCATCGGA |
| *NEU1* | CAGCACATCCAGAGTTCCGAGT | TGTCTCTTTCCGCCATGAGGT |
| *PIP4P1* | GTTCGATGCCCCTGTAACTGTC | CCCAGGTTGATGATTCTTTTGC |
| *SCPEP1* | GATCTCCCCTGTTGATTCGGT | AGCCCCTTATTTACGGCATTC |
| *TFEB* | CCAGAAGCGAGAGCTCACAGAT | TGTGATTGTCTTTCTTCTGCCG |
